# UBE2C promotes leptomeningeal dissemination and is a therapeutic target in brain metastatic disease

**DOI:** 10.1101/2022.10.04.510674

**Authors:** Eunice Paisana, Rita Cascão, Carlos Custódia, Nan Qin, Daniel Picard, David Pauck, Tânia Carvalho, Pedro Ruivo, Clara Barreto, Delfim Doutel, José Cabeçadas, Rafael Roque, José Pimentel, José Miguéns, Marc Remke, João T. Barata, Claudia C. Faria

## Abstract

Despite current improvements in systemic cancer treatment, brain metastases (BM) remain incurable, and there is an unmet clinical need for effective targeted therapies. Here, we sought common molecular events in brain metastatic disease. RNA sequencing of thirty human BM identified the upregulation of *UBE2C*, a gene that ensures the correct transition from metaphase to anaphase, across different primary tumor origins. Tissue microarray analysis of an independent BM patient cohort revealed that high expression of UBE2C was associated with decreased survival. UBE2C-driven orthotopic mouse models developed extensive leptomeningeal dissemination, likely due to increased migration and invasion. Early cancer treatment with dactolisib (dual PI3K/mTOR inhibitor) prevented the development of UBE2C-induced leptomeningeal metastases. Our findings reveal UBE2C as a key player in the development of metastatic brain disease and highlight PI3K/mTOR inhibition as a promising anticancer therapy to prevent late-stage metastatic brain cancer.

## INTRODUCTION

Dissemination of cancer cells to the brain is a frequent and late-stage complication of many systemic cancers. Patients with brain metastatic disease have a poor prognosis, with a median survival between 3 to 11 months^1,2^. Lung cancer (40-60%), breast cancer (15-30%), and melanoma (5-15%) are the most common primary malignancies that disseminate to the brain^2-5^. Autopsy studies showed that around 20 to 26% of all cancer patients develop brain metastases (BM) but it is thought that the incidence may be higher and increasing^3,5,6^. Advanced methods of BM diagnosis and more effective treatments for extracranial disease led to an improvement in patients’ survival, thus contributing to an increase in BM incidence. The standard of care treatment for BM includes surgical resection, whole-brain radiotherapy (WBRT), and stereotactic radiosurgery (SRS)^7^. Nonetheless, brain metastatic disease remains incurable^3^. Furthermore, the most aggressive form of this disease is characterized by coating of 18the leptomeninges by cancer cells (leptomeningeal dissemination), and poor response to intensive chemotherapeutic regimens, including intrathecal therapy^8^. Currently, molecular-targeted therapies in use for primary tumors lack good therapeutic responses in metastatic tumors located in the central nervous system (CNS). This might be explained either by the limited crossing of drugs through the blood-brain barrier (BBB)^9^ or by genetic differences between primary tumors and BM^10^.

Understanding the molecular mechanisms underlying the dissemination of cancer cells into the brain will open the possibility of identifying targets to be used in the development of novel therapies. Previous attempts to study the mechanisms of cancer dissemination to the brain focused mainly on analyzing genes differentially expressed between the most common primary tumors (lung and breast cancer) and their BM. The metastatic compartment often exhibits a distinct genetic profile from the primary tumor^10^. Specific molecular targets have been found to play a role in the metastatic process to the brain, namely HER2, FOXC1, LEF1 and HOXB9 in lung cancer, COX2, HBEGF, and ST6GALNAC5 in breast cancer, and plasminnogen activator (PA) inhibitory serpins in both these tumors^11-13^.

We hypothesized that brain metastatic tumors from diverse histological types, share common genetic events which promote cancer cell dissemination and colonization of the brain, and these molecular alterations can be used as therapeutic targets. We performed RNA sequencing in a discovery cohort of thirty BM samples from patients diagnosed with various primary tumors and identified UBE2C as being differentially expressed. UBE2C is a ubiquitin-conjugating enzyme that functions together with anaphase-promoting complex/cyclosome (APC/C) involved in the cell cycle, ensuring a correct transition from metaphase to anaphase^14,15^. High expression of UBE2C has been found in tumor tissues, and it was associated with a worse prognosis in primary cancers^16,17^. We analyzed an independent cohort of 89 BM from different primary origins and observed that patients with higher expression of UBE2C had a significantly worse prognosis. In orthotopic mouse models of BM, UBE2C promoted leptomeningeal dissemination and decreased survival, possibly due to an increase in cancer cell migration and invasion. Importantly, early treatment with a PI3K/mTOR inhibitor (dactolisib) prevented the UBE2C-induced leptomeningeal dissemination in these models, revealing a promising therapeutic avenue in advanced metastatic cancer prevention.

## RESULTS

### *UBE2C* is upregulated in human BM

A discovery cohort from the neurosurgery department at Hospital de Santa Maria from Centro Hospitalar Universitário Lisboa Norte (HSM-CHULN) comprising thirty human BM from patients diagnosed with different primary tumor origins was analyzed using RNA sequencing. In this cohort, lung cancer was the most frequent primary tumor, followed by breast, uterus and colon cancer (**Fig. 1A**). Tumor samples were collected consecutively, and therefore, the representation of each primary tumor origin reflects the frequency in the neurosurgical series. The transcriptomic analysis included publicly available RNA sequencing datasets of normal tissues (**Fig. 1B**) and microarray datasets of normal tissue and primary tumors (**Fig. S1A**), matching the BM cohort. Genes upregulated exclusively in BM samples were identified (n=4514 genes, **Fig. 1C**). From this list, we selected the top 20 upregulated genes based on *p*-value ≤0.05 and positive or negative fold change ≥1.7 (**Fig. 1D** and **Fig. S1B**). The expression of candidate genes was then checked for directionality in all tumor types and the most promising genes were further investigated regarding their clinical relevance. *UBE2C* was found to be highly expressed in BM samples from diverse primary tumor origins when compared to normal tissues (**Fig. 1E**). Importantly, among the top 5 candidate genes, *UBE2C* was the most frequently associated with poor prognosis in publicly available clinical datasets from diverse cancer types, including metastatic melanoma (**Fig. 1F**).

**Figure 1.**
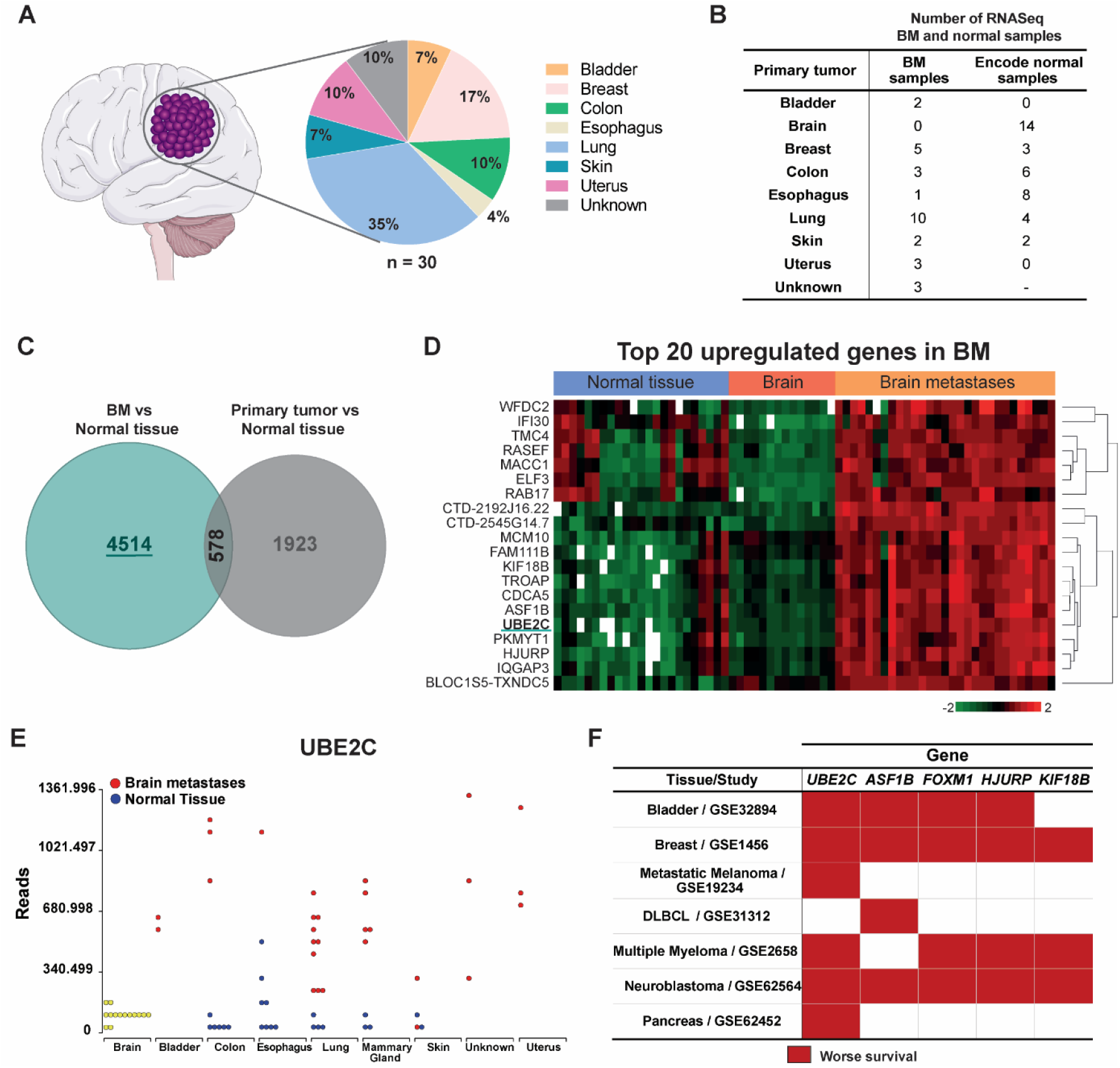
UBE2C is highly expressed in human BM. **A**. Percentage of BM included in the RNA sequencing analysis, according to primary tumor origin (n=30, discovery cohort from HSM-CHULN). **B**. Datasets included in the RNA sequencing (RNASeq) bioinformatic analysis, including data from normal tissue samples, from publicly available datasets. **C**. Venn diagram analyzing the differentially expressed genes in BM and primary tumor samples, against genes from normal tissue. **D**. Heatmap representing the top 20 upregulated genes in our discovery cohort when compared to publicly available data from normal brain and other normal tissues. **E**. Comparison of RNA UBE2C levels in BM and normal tissues. **F**. Top 5 upregulated genes in BM and their association with the overall survival (OS) of cancer patients, using datasets from several studies with publicly available RNAseq data: GSE32894: Hoglund, 2015^18^; GSE1456: Bergh, 2005^19^; GSE19234: Bhardwaj^20^, 2009; GSE31312: International DLBCL Rituximab-CHOP Consortium^21^; GSE2658: Hanamura, 2006^22^; GSE62564: Tong, 2014^23^; GSE62452: Hussain, 2016^24^. DLBCL: Diffuse large B-cell lymphoma. Red color represents high gene expression significantly associated with decreased OS.

### Patients with high expression of UBE2C in BM have poorer prognosis

We validated the clinical relevance of UBE2C by immunohistochemistry (IHC) in tissue microarrays (TMAs), using an independent cohort of 89 patients with BM from different primary tumor origins (**Fig. 2A-B**). Samples from patients with primary brain tumors (glioblastoma) were used as control. Patients had a median age of 62 years (28-90 years) (**Fig. S2A**), with a similar frequency of male and female patients (52% and 48%, respectively) (**Fig. S2B**). The median survival of these patients since the diagnosis of BM was 8 months (**Fig. S2C**), with different survival rates depending on the primary tumor origin (**Fig. S2D-E**). High expression of UBE2C was correlated with a worse prognosis (median survival of 7 months), while patients with low UBE2C expression exhibited a median survival of 12 months (*p-*value=0.04) (**Fig. 2C**). This finding was not due to a higher proliferative index in tumor cells (**Fig. 2D-E**), nor dependent on the primary tumor (**Fig. S2F-J**). Using a scoring system combining intensity and frequency of UBE2C staining (**Fig. 2F**), we have observed that most BM samples (55%) have high levels of UBE2C (Score III and IV), in contrast with glioblastomas in which 70% of the samples have low expression of this protein (**Fig. 2G**). This suggests that high UBE2C expression is a feature of brain metastatic tumors.

**Figure 2.**
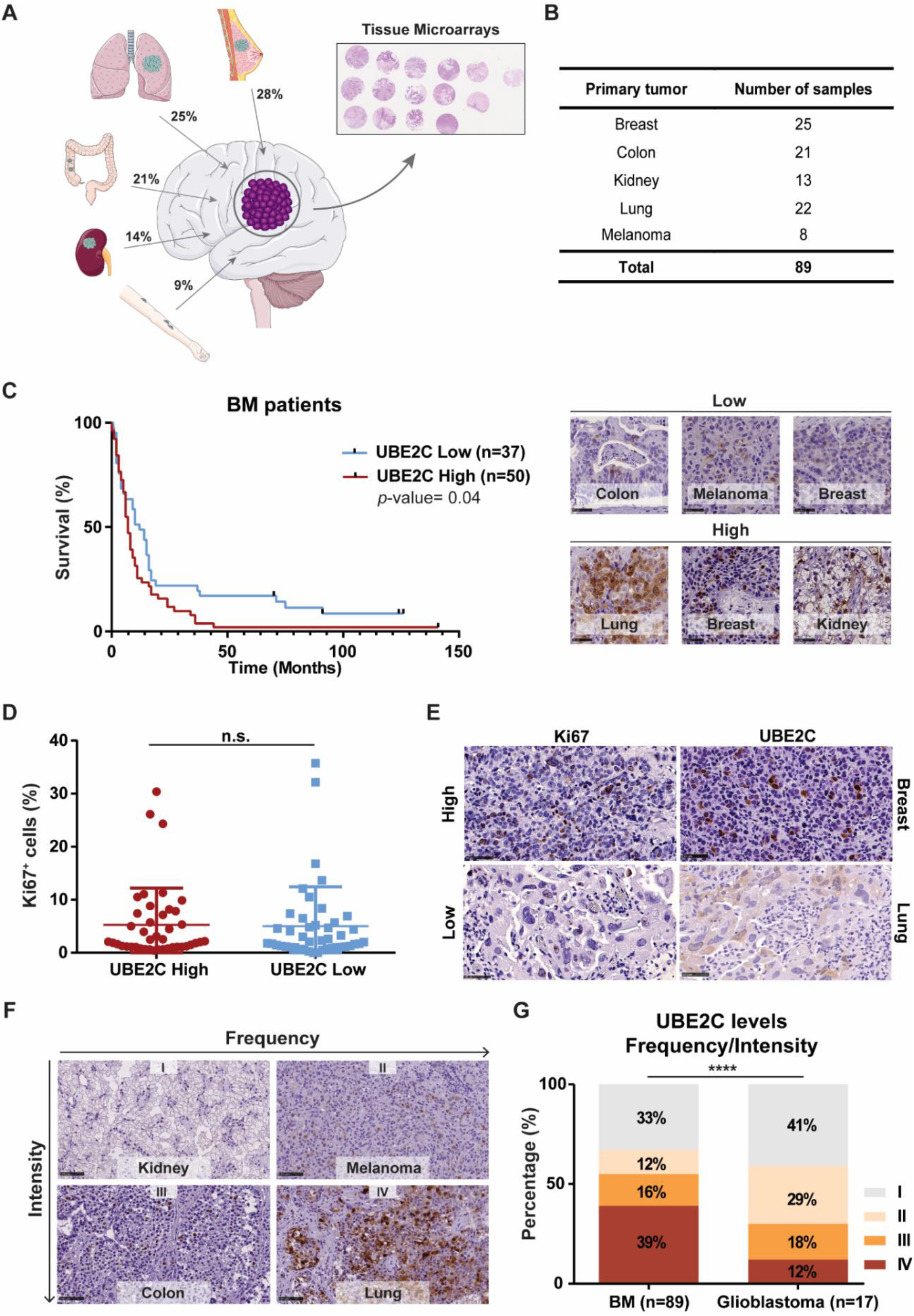
High expression of UBE2C is associated with worse survival in patients with BM. **A**. Percentage of BM by tumor type included in the tissue microarrays (TMA). **B**. Number of BM samples by tumor origin in the TMA (independent validation cohort, N=89). **C**. Kaplan-Meier analysis of patients’ survival according to UBE2C protein levels and representative images of the IHC intensity score used (high and low). Differences were considered statistically significant for *p-*values≤0.05, according to the Log-rank (Mantel-Cox) test. **D**. Comparison of UBE2C and Ki67 staining in patients with BM. Data is expressed as median with interquartile range. Mann–Whitney test. **E**. Representative images of IHC staining for UBE2C (scored as high or low) and Ki67 (assessed using ImageJ); Scale bar: 50μm. **F**. IHC score including intensity and frequency of the UBE2C staining in tumor tissue: I – low/low; II – low/high; III – high/low; III – high/high, respectively; Scale bar: 100μm. **G**. UBE2C expression was compared between BM and brain primary tumors (glioblastoma); \*\*\*\**p*-value< 0.0001, Chi-square (and Fisher’s exact) test.

### UBE2C promotes cancer cell migration and invasion *in vitro*

To characterize the role of UBE2C in the context of brain metastatic disease we used two cancer cell lines, MDA (breast cancer) and A549 (lung cancer). Both cell lines were engineered to stably overexpress *UBE2C* (**Fig. 3A-B**). To study the effect of UBE2C in cancer cell migration, we performed real-time cell analyses using the xCELLigence system (**Fig. 3C**). The migration rate of MDA and A549 cells was significantly increased in UBE2C overexpressing cells (**Fig. 3D-E**). This effect was not due to an increase in proliferation either in short-term (**Fig. S3A-B**) or long-term assays (**Fig. 3F-H** and **Fig. S3C-E**), as assessed by MTS and colony formation assays, respectively. Additionally, UBE2C increased the invasion ability of MDA cells in matrigel-coated transwells (**Fig. 3I-J**).

**Figure 3.**
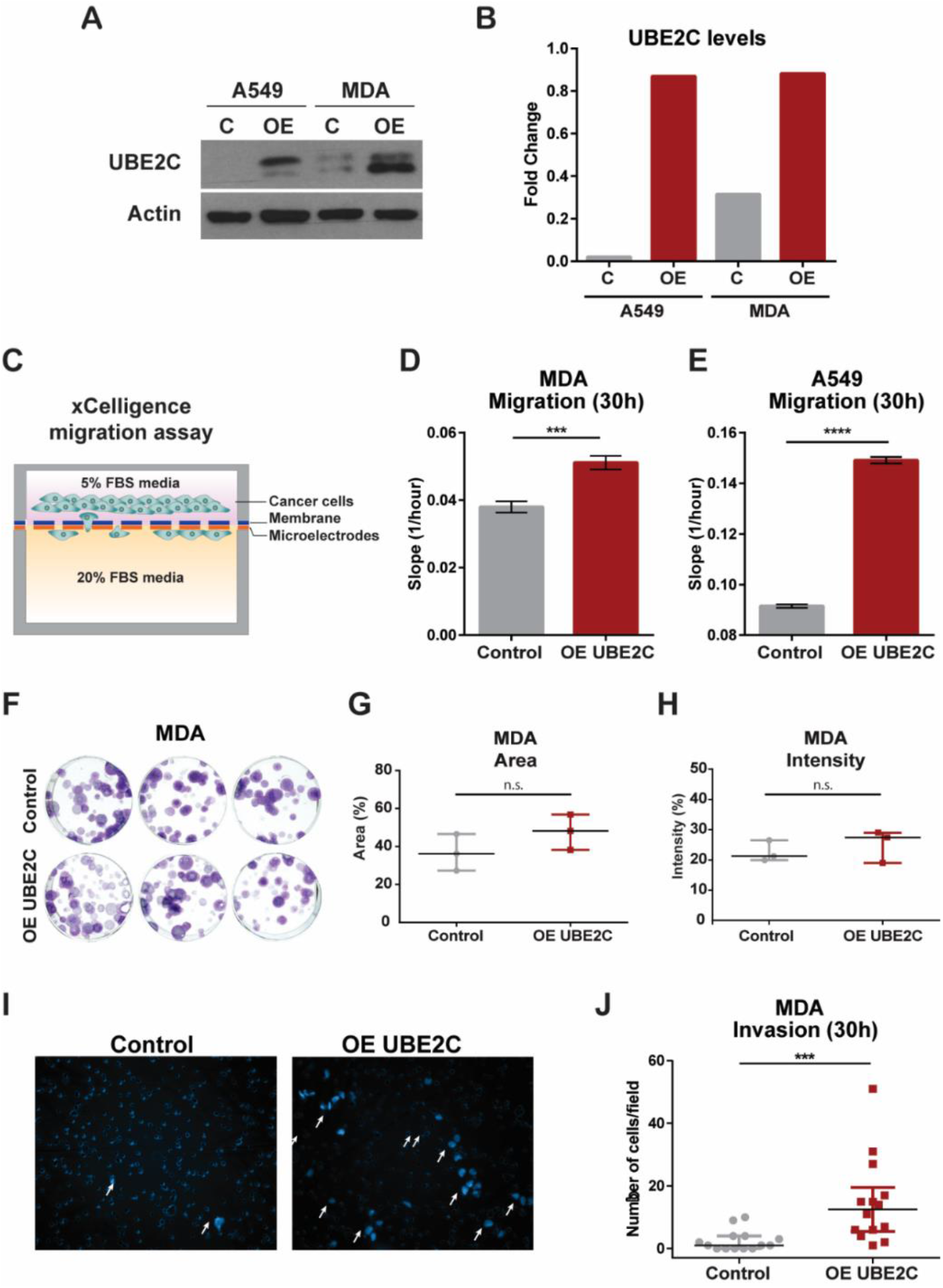
Overexpression of *UBE2C* increases cancer cell migration and invasion. **A**. Western Blot of UBE2C levels in MDA and A549 cell lines modulated for overexpression (OE) of *UBE2C* and **B**. its densitometry analysis. **C**. Schematic representation of migration assay using CIM-Plates in the xCELLigence system. Migration ability of (**D**) MDA and (**E**) A549 cells with OE of *UBE2C* was evaluated using the xCELLigence system at 30h and compared with control cells (empty vector); \*\*\*\**p*-value< 0.0001, \*\*\**p*-value= 0.001, unpaired t-test. Data represented as mean with SD. **F**. Pictures of colony formation assay (CFA) performed in MDA OE *UBE2C* cells (100 cells/well) after 2 weeks, and (**G**) quantification of the area; *p*-value=0.23 and (**H**) intensity *p*-value=0.52 of the formed colonies. Mann-Whitney test. Data is presented as median and interquartile range. **I**. Representative images and (**J**) quantification of the invasion capacity of MDA cells with OE *UBE2C* using matrigel-coated transwells at 30h and compared with control cells (empty vector); \*\*\**p*-value= 0.0002, Mann-Whitney test. Data is presented as median with interquartile range.

### UBE2C drives leptomeningeal dissemination and associates with worse survival *in vivo*

To assess the *in vivo* role of UBE2C in BM, we performed intracranial injections of MDA cells in NSG mice (**Fig. 4A**). Animals injected with *UBE2C*-overexpressing cells showed decreased survival (**Fig. 4B**, *p*-value=0.05), a phenotype that associated with more aggressive disease characterized by leptomeningeal dissemination. Although no differences were found in leptomeningeal dissemination in the brain (**Fig. S4A**), we observed a significant increase in spine dissemination both *in vivo* (**Fig. 4C-D**) and *ex vivo* (**Fig. 4E-G**), in animals with *UBE2C*-overexpressing cells. We also established an MDA cell line with *UBE2C* knock down (KD) using the CRISPR system (**Fig. S4B**). There were no differences in survival (**Fig. S4C**) or dissemination to the brain meninges (**Fig. S4D**) in orthotopic models transplanted with *UBE2C* KD or control cells. However, we found evidence of decreased dissemination of cells with *UBE2C* silencing to the spinal cord meninges (**Fig. S4E**). Thus, *UBE2C* drives a more aggressive disease phenotype with leptomeningeal dissemination and decreased survival in orthotopic models of BM.

**Figure 4.**
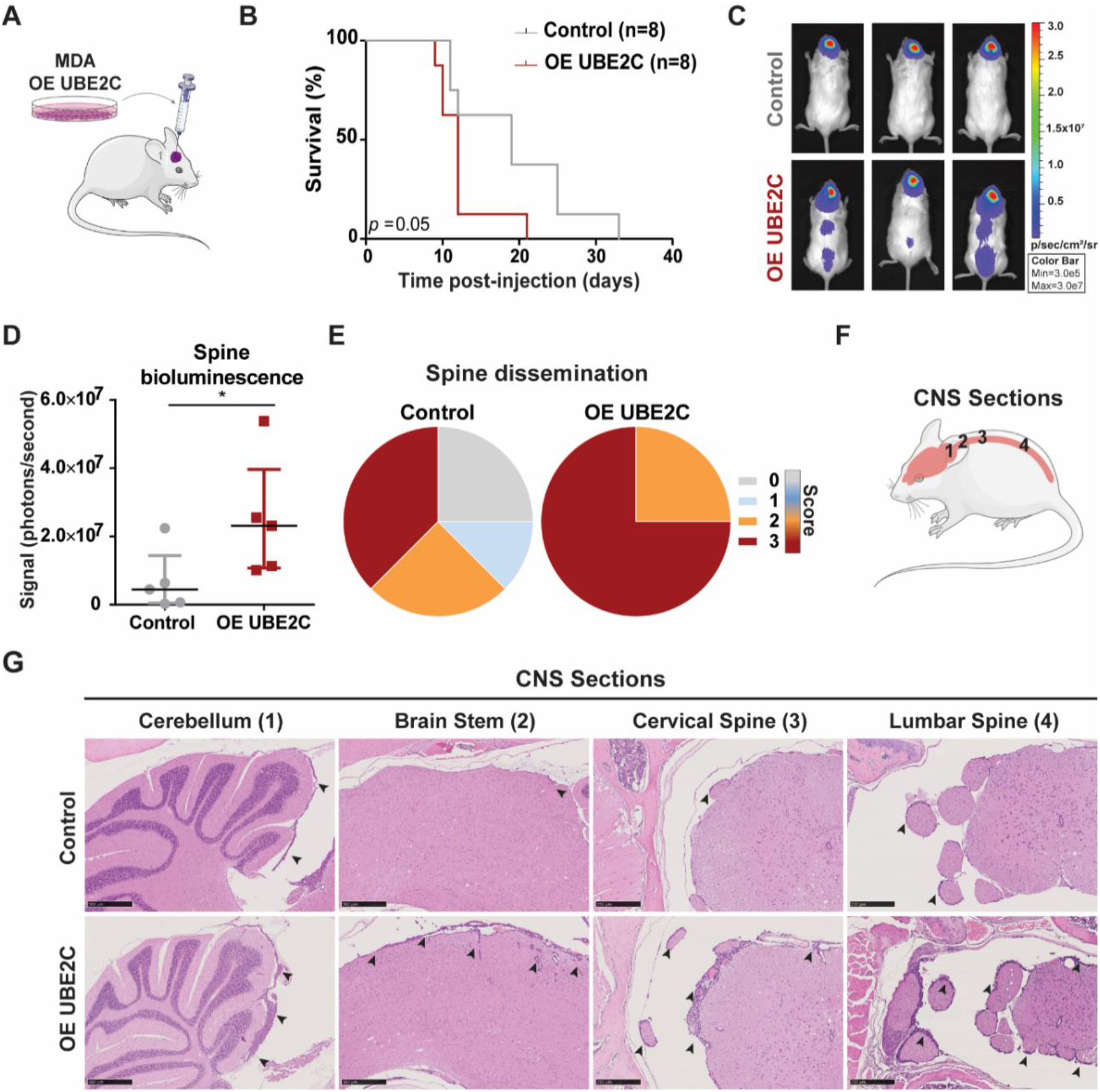
*UBE2C* decreases survival and mediates leptomeningeal dissemination driving an aggressive disease phenotype *in vivo*. **A**. Intracranial injection of NSG mice (n=8/group) with MDA cell line, control and OE *UBE2C;* **B**. and the survival analysis of both groups, \**p-*value=0.05, Kaplan-Meier test. **C**. Representative images of the whole-body bioluminescence imaging of control and OE *UBE2C* mice at day 9 post-injection, and (**D**) quantification of the bioluminescence signal in the spine; \**p*-value=0.03, Mann-Whitney test. Data represented as median with interquartile range. **E**. Spine leptomeningeal dissemination in animals injected with control and OE *UBE2C* cells. Score used to assess the leptomeningeal dissemination: 0-negative; 1-mild; 2-moderate; 3-marked. **F**. CNS sections analyzed by histopathology, and (**G**) representative images of these sections with H&E staining (5x; scale bar: 250μm) from mice injected with control and OE *UBE2C* cells. Black arrows indicate leptomeningeal dissemination.

### Targeting the PI3K/mTOR pathway *in vitro* decreases cancer cell proliferation and downregulates UBE2C

Due to the lack of effective treatment options for patients with BM and the inexistence of specific therapies targeting UBE2C, we performed a high-throughput drug screening to identify compounds with efficacy in treating UBE2C-driven BM. We used a chemical library of 650 compounds, FDA approved or in phase 3 or 4 clinical trials, to test cell lines with overexpression and knock-down of UBE2C (**Fig. S5A**). We have selected two candidates for *in vitro* validation which targeted cells with high expression levels of UBE2C: dactolisib (PI3K/mTOR inhibitor) and Genz644282 (Topoisomerase I inhibitor) (**Fig. S5B-C**). Both compounds proved to be highly effective inhibiting cancer cell proliferation (**Fig. 5A-B** and **S5D-E**).

**Figure 5.**
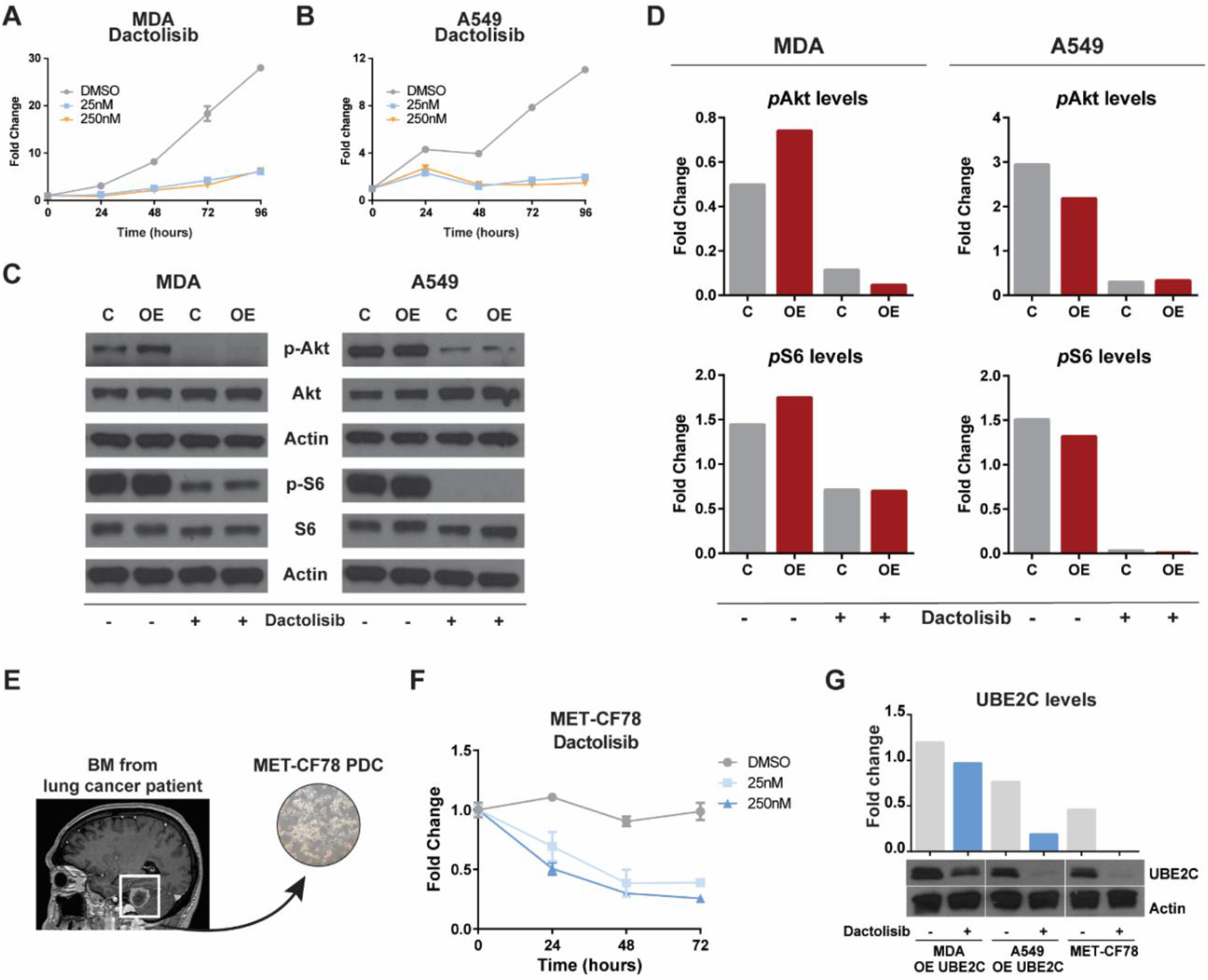
Downregulation of PI3K/mTOR pathway inhibits cancer cell proliferation and decreases UBE2C levels. MTS assays using 25nM and 250nM concentrations of dactolisib to assess proliferation in (**A**) MDA and (**B**) A459 cell lines with OE of *UBE2C*. **C**. Evaluation of the effect of dactolisib in PI3K/mTOR pathway (*p*-Akt and *p*-S6 levels) by western blot using MDA and A549 cells with *UBE2C* overexpression (OE) in comparison with control cells (**D**). **E**. MET-CF78 PDC derived from a BM of lung cancer patient. **F**. Inhibition of proliferation by dactolisib in patient-derived cultures from a lung cancer BM (MET-CF78). **G**. UBE2C levels assessed by western blot in cancer cell lines treated with dactolisib (250nM) for 24h and its densitometry analysis.

Single-cell RNA sequencing data analysis of BM from patients with different primary tumors^25^ showed that metastatic tumor cells with high UBE2C expression levels (**Fig. S5F**) also presented high levels of MTOR1 (**Fig. S5G**), with a positive correlation between the mRNA levels of both genes (**Fig. S5H**). Additionally, the PI3K/mTOR pathway has been implicated in brain metastatic cancer^10^. Therefore, we decided to further evaluate the efficacy of inhibition of PI3K/mTOR signaling and examine its interplay with UBE2C. Dactolisib effectively inhibited the PI3K/mTOR pathway in breast and lung cancer cell lines *in vitro*, through downregulation of pAkt and pS6, respectively (**Fig. 5C-D**). Furthermore, we have also used a patient-derived culture (PDC), MET-CF78, isolated from a BM derived from a lung cancer patient with constitutive expression of UBE2C (**Fig. 5E**)^26^. We observed inhibition of MET-CF78 proliferation by dactolisib in a dose-dependent manner (**Fig. 5F**). Interestingly, the targeting of PI3K/mTOR signaling pathway by dactolisib decreased the UBE2C levels in all cancer cell lines (**Fig. 5G**).

### Dactolisib prevents leptomeningeal dissemination *in vivo*

We then asked whether the aggressive metastatic phenotype induced by UBE2C *in vivo* could be prevented by early treatment with dactolisib, a compound known to cross the BBB^27^. A preclinical therapeutic protocol was designed to early treat UBE2C-driven orthotopic mouse models of BM with dactolisib by oral gavage. Treatment was initiated on day 4 post-injection, and it was given daily for 10 days, with a wash-out period of two days (**Fig. 6A**). There were no differences in weight between dactolisib-treated and untreated animals (**Fig. S6A**). Strikingly, although dactolisib was not able to significantly reduce brain tumor size (**Fig. S6B**), it was effective in preventing leptomeningeal dissemination both in brain and spine (**Fig. 6C-F**), reversing the aggressive phenotype induced by *UBE2C*-overexpressing cancer cells. Dactolisib-treated brain tumors exhibited a significant decrease in the number of cancer cells expressing high levels of UBE2C (**Fig. 6G-H**).

**Figure 6.**
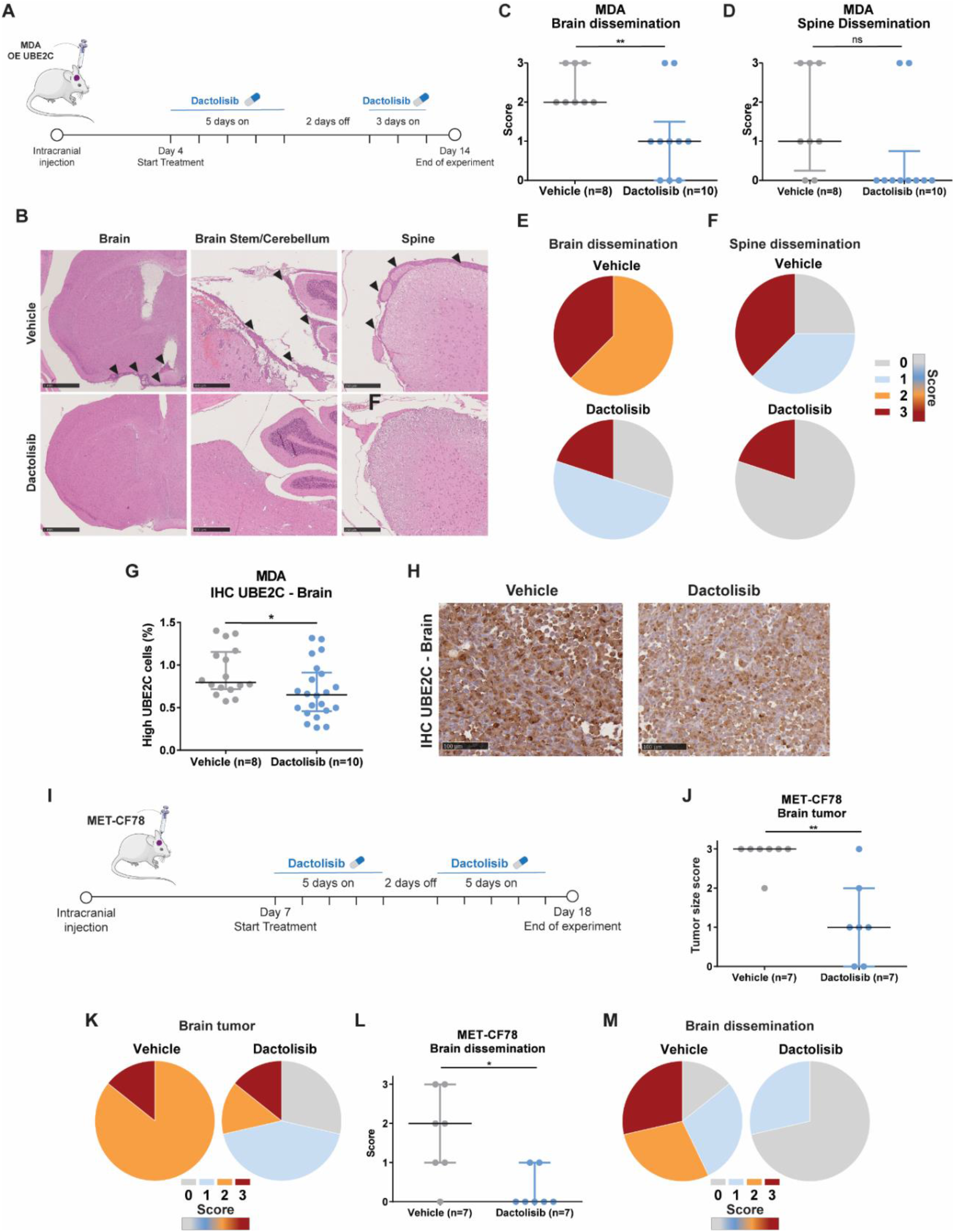
PI3K/mTOR dual inhibitor dactolisib prevents leptomeningeal dissemination in *UBE2C*-driven mouse models of BM. **A**. Treatment protocol in orthotopic xenografts using breast cancer cells (MDA) with overexpression of *UBE2C*. Animals were treated daily from day 4 to day 14 post-injection with dactolisib (30mg/kg; n=10) or vehicle (NMP and PEG300, 10%-90%, v-v; n=8), with a wash-out period of 2 days. **B**. Representative images of leptomeningeal dissemination in the control group and absence of dissemination in the dactolisib-treated group. Brain (scale: 1mm), brain stem/cerebellum (scale: 500μm) and spine (scale: 250 μm). **C**. Histopathological scoring of leptomeningeal dissemination in the brain (\*\**p*-value=0.0083, t-test, data represented as mean and SD) and **D**. in the spine (*p*-value=0.07, Mann-Whitney test, data represented as median and interquartile range). **E**. Percentage of brain dissemination in vehicle (2: 62,5%, 3: 37,5%) and treatment (0: 30%, 1: 50%, 3: 20%) groups. **F**. Percentage of spine dissemination in vehicle (0: 25%, 1: 37,5%, 3: 37,5%) and treatment (0: 80%, 3: 20%) groups. **G**. Expression of UBE2C by IHC staining, in brain tumor samples upon treatment with dactolisib. \**p*-value=0.03, t-test. Data represented as mean with SD. **H**. Representative images of UBE2C staining in brain tumor sections. **I**. Treatment protocol in patient-derived xenografts of lung cancer BM (MET-CF78) (n=7/group). Animals were treated daily from day 7 to day 17 post-injection with dactolisib or vehicle, with a wash-out period of 2 days; Scale bar: 100μm. **J**. Histopathological scoring of the brain tumor size (0: no tumor; 1: minimal to mild; 2: moderate; 3: marked); \*\**p*-value=0.009; Mann-Whitney test. Data represented as median with interquartile range, and **K**. brain tumor size score in vehicle (2: 14,286%, 3: 85,714%) and dactolisib (0: 28,571%, 1: 42,857%, 2: 14,286%, 3: 14,286%) treated mice. **L**. Histopathological scoring of brain dissemination as described above; \**p*-value=0.026, Mann-Whitney test; data represented as median with interquartile range, and **M**. percentage of brain dissemination in vehicle (0: 14,286%, 1: 28,571%, 2: 28,571%, 3: 28,571%) and treatment (0: 81,429%, 1: 28,571%) groups. Score used to assess the leptomeningeal dissemination: 0-negative; 1-mild; 2-moderate; 3-marked

In the patient-derived xenograft of the lung cancer BM (MET-CF78), early preclinical treatment with oral dactolisib (**Fig. 6I**) had no differences in spine dissemination (**Fig. S6C**) and there was no impact of the treatment in the weight of the animals (**Fig. S6D**). However, dactolisib significantly reduced both brain tumor size (**Fig. 6J-K**) and leptomeningeal dissemination to the brain (**Fig. 6L-M**).

In order to validate that the effect of dactolisib *in vivo* was, at least in part, *UBE2C*-dependent, we developed orthotopic mouse models using MDA cells with KD of *UBE2C* (n=7 mice/group). There were no differences in the weight of treated animals compared to the untreated (**Fig. S6E**). In UBE2C-depleted tumors, dactolisib did not significantly impact brain tumor size (**Fig. S6F**) or CNS dissemination (**Fig. S6G-H**).

## DISCUSSION

Despite significant therapeutic advances in the treatment of primary cancers, BM remain a major clinical hurdle. We began addressing this challenge by interrogating BM from diverse primary cancers using RNA sequencing coupled with functional approaches, in both human and mouse models. Focusing on BM from diverse primary tumors, we were able to identify common genetic events leading to cancer cell dissemination and/or colonization into the brain. Simultaneously, these molecular targets constitute novel targets for therapy.

We identified UBE2C as a differentially expressed gene in human BM and showed that high levels of UBE2C are associated with shorter survival in cancer patients with brain metastatic disease. UBE2C was first described by Okamoto as an oncogene, overexpressed in primary tumors and cancer cell lines, when comparing with normal tissues^16^. It has also been shown that UBE2C correlates with higher tumor grades. In lung cancer patients, UBE2C was associated with poorer survival^28^ and with tumor progression, as a consequence of autophagy inhibition and increased tumor invasiveness^29^. In breast cancer, UBE2C was identified as a prognostic marker^30^ and associated with cancer grade progression^31^ and drug resistance^32^. Similar observations were made in colon, ovarian, and bladder cancers, among other types of tumors^33-36^. Interestingly, UBE2C was associated with BM in a recent publication, where single-cell RNA sequencing of BM from multiple cancers led to the identification of two subgroups, being UBE2C one of the signature genes in the proliferative group^25^.

UBE2C has been described as a promoter of migration and invasion of tumor cells in endometrial cancer models, inducing epithelial to mesenchymal transition (EMT), via downregulation of p53 levels^37^. Similar effects were also observed in gastric^38^ and hepatocellular cancers^39^. We observed that orthotopic mouse models injected with cancer cells overexpressing UBE2C developed a more aggressive disease phenotype, with leptomeningeal dissemination in the brain and spinal cord. This phenomenon may be explained by the increase in cancer cell migration and invasion induced by overexpression of this gene, as observed *in vitro*.

In cancer patients, leptomeningeal dissemination can occur in approximately 5-15% of the cases^8^, having a dismal prognosis with a median survival between 2 to 4 months^8,40-42^. Very little is known about the molecular biology of this advanced stage of metastatic cancer, making it exceedingly difficult to treat. Aggressive therapies including intrathecal chemotherapy and whole brain radiation have failed to alter the natural history of leptomeningeal dissemination. Recent phase 2 studies have reported clinical benefit of patients treated with EGRF inhibitors and immunotherapy^43,44^, although the duration of responses in leptomeningeal disease was limited in time.

Since late-stage CNS dissemination seems very difficult to target, the best chance for an improved survival in these patients would be to prevent cancer cells from disseminating into the leptomeninges. Our observations revealed that early treatment with oral dactolisib (a dual PI3K/mTOR inhibitor) prevented the development of leptomeningeal dissemination in orthotopic mouse models driven by UBE2C. Moreover, dactolisib effectively reduced dissemination and brain tumor size in a patient-derived xenograft from a lung cancer BM. In both *in vivo* models, the dual PI3K/mTOR inhibitor was well tolerated and did not induce toxicity. Interestingly, mutations in PI3K/Akt/mTOR signaling have been found in BM, but not in primary tumors^10^. Also, activation of this signaling pathway has already been associated with leptomeningeal dissemination in melanoma patients^45^. We have shown that PI3K/mTOR pathway inhibition with dactolisib decreased cancer migration and invasion, and downregulated UBE2C both *in vitro* and *in vivo*. The connection between PI3K/Akt/mTOR pathway and UBE2C has been previously reported in gastric cancer cells, where UBE2C led to the activation of AURKA and consequent EMT by the decrease in p-AKT1 levels^38^. Furthermore, the mTOR inhibitor CCI-779 decreased the levels of UBE2C in castration-resistant prostate cancer, by disrupting its transcription and blocking UBE2C-dependent invasion^46^. We believe dactolisib targets key steps in the metastatic cascade (mobilization of cancer cells from the primary tumor and invasion) and, therefore, may be considered as a novel therapeutic strategy to prevent brain metastatic disease.

Further studies are warranted to better explore the mechanisms of UBE2C-driven dissemination to the brain and to dissect the cellular interplay between cancer, stroma and immune cells, mediated by UBE2C. Moreover, the limited size of our BM patients’ cohort, does not allow to extrapolate our results for each cancer type and molecular subtype. The comparison of UBE2C expression levels in BM with matched primary tumors or extracranial metastases would also add layers of information on UBE2C specificity.

This notwithstanding, we have showed that UBE2C is a relevant player in brain metastatic disease, especially relevant for leptomeningeal dissemination, and that it can be used as a prognostic marker in cancer patients with this condition. Furthermore, we demonstrated that UBE2C-induced leptomeningeal dissemination can be prevented *in vivo* by targeting the PI3K/mTOR pathway (**Fig. 7**). The results from our study may prompt the advance of PI3K/mTOR inhibitors into clinical trials for patients with advanced metastatic cancer of the CNS from multiple primary origins.

**Figure 7.**
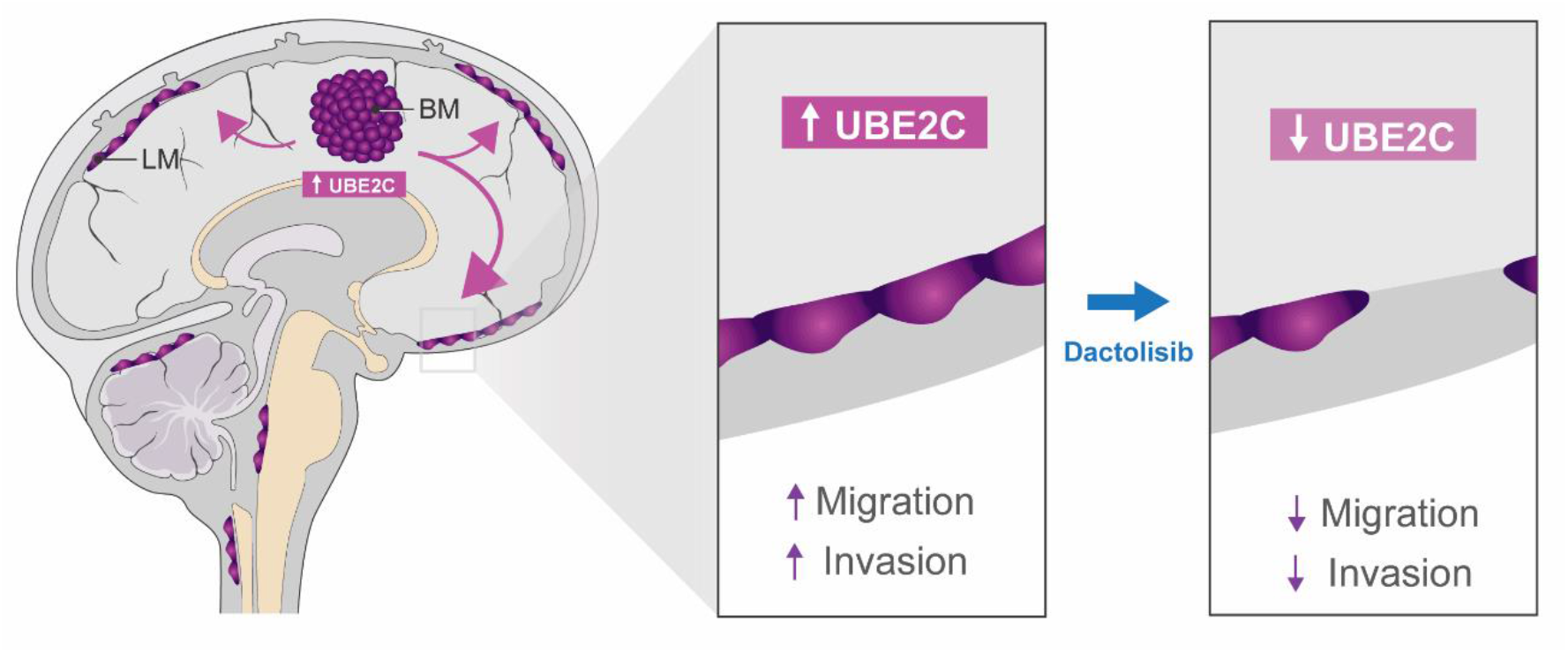
Working Model. High levels of UBE2C in BM increase migration and invasion capacity of cancer cells and promote leptomeningeal dissemination. Treatment with dactolisib (PI3K/mTOR pathway inhibitor) prevents this late-stage complication of cancer patients.

## METHODS

### Patients’ samples

BM samples and detailed clinical data were collected from 30 patients with diverse primary tumors (lung, breast, uterus, bladder, colon, esophagus and melanoma), submitted to surgical resection in the Department of Neurosurgery at HSM-CHULN. Surgical specimens not needed for diagnostic purposes were snap frozen in liquid nitrogen and stored at Biobanco-iMM CAML (biobank of Lisbon Academic Medical Center, Lisbon, Portugal) within an hour after surgery. BM samples were collected in accordance with the Ethics board from Hospital de Santa Maria (Ref^a^. Nº 367/18 and Ref^a^. Nº 346/20) and a written informed consent was obtained from all patients, prior to study participation. Samples were then requested from Biobanco-iMM for RNA sequencing analysis, as described below.

### RNA isolation from BM specimens

Snap frozen samples of BM were used in the RNA sequencing analysis. Total RNA was isolated from BM tissue (500ng) using Trizol™ reagent (Invitrogen-Thermo Fisher Scientific, Waltham, Massachusetts, USA, Cat#15596026) according to the manufacturer’s recommendations. Following isolation, RNA was stored in RNase/DNase-free water at −80°C. The quantity and quality of the isolated RNA was assessed by NanoDrop™ (Thermo Fisher Scientific, Waltham, Massachusetts, USA). Samples were excluded if the yield did not reach the minimum requirement of 1000ng or RIN>6 (RNA Integrity Number).

### RNA sequencing analysis

Total RNA isolated from BM was processed using the TruSeq RNA Sample Preparation v2 kit (low-throughput protocol; Illumina, San Diego, CA, USA) to prepare the barcoded libraries. Libraries were validated and quantified using either DNA 1000 or high-sensitivity chips on a Bioanalyzer (Agilent, Santa Clara, CA, USA). 7.5 pM denatured libraries were input into cBot (Illumina), followed by deep sequencing using HiSeq 2500 (Illumina) for 101 cycles, with an additional seven cycles for index reading. For normal tissue samples, we downloaded the call sets from the ENCODE portal^47^ (https://www.encodeproject.org/) for tissues matching the tissue of origin of the BM.

Fastq files were imported into Partek Flow (Partek Incorporated, St. Louis, MO, USA). Quality analysis and quality control were performed on all reads to assess read quality and to determine the amount of trimming required (both ends: 13 bases 5’ and 1 base 3’). Trimmed reads were aligned against the hg38 genome using the STAR v2.4.1d aligner. Unaligned reads were further processed using Bowtie 2 v2.2.5 aligner. Finally, aligned reads were combined before quantifying the expression against the ENSEMBL (release 84) database using the Partek Expectation-Maximization algorithm. Partek Flow default settings were used in all analyses. Files were then processed using Partek Genomic Suite (Partek Incorporated, St. Louis, MO, USA). Genes were filtered for expression values ≤1 and 3 or more missing values, remaining genes were then log2 transformed.

### Microarray datasets used in the bioinformatic analysis

Microarray datasets were also used in the analysis of RNA sequencing. GSE2109 and GSE7307 datasets were downloaded from GEO DataSets (https://www.ncbi.nlm.nih.gov/gds) and processed using Partek Genomic Suite. Files were imported into Partek Genomic Suites and normalized using the RMA method.

### Tissue Microarrays of human BM

Tissue microarrays (TMAs) were previously built in the Neuropathology lab of HSM-CHULN and kindly provided by Pedro Pereira and Professor José Pimentel. The TMA included samples of BM collected from patients with primary tumors from diverse origins (**Fig. 2A**). UBE2C protein levels were assessed by immunohistochemical (IHC) staining with UBE2C antibody (Boston Biochem, Cat# A650) or Ki-67 (D2H10) (Cell Signaling Technology, Leiden, The Netherlands, Cat# 9027) using 5μm sections of the formalin fixed, paraffin embedded (FFPE) TMA. FFPE sections were incubated with the primary antibody and with EnVision+ (Dako, Glostrup, Denmark). Color was developed in solution containing diaminobenzadine-tetrahydrochloride (Sigma, Missouri, USA), 0.5% H_2_O_2_ in phosphate-buffered saline buffer (pH 7.6). Slides were counterstained with hematoxylin and mounted. IHC evaluation was performed blindly by 2 independent researchers and a specialized pathologist. For UBE2C we used a semi-quantitative score of intensity (low or high staining intensity) and frequency (low: 0-49% staining or high: 50-100% staining). For Ki67, an automated software was used to quantify the percentage of positive nuclei (ImmunoRatio). Images were acquired using a NanoZoomer SQ slide scanner (Hamamatsu Photonics, Hamamatsu City, Japan) with 20x magnification (0.46μm resolution).

### Cell culture of cell lines and induction of brain tropism *in vivo*

Human cell lines MDA-MB-231, A549 and HCT116 were maintained in the appropriate media and passaged up to 15 times. MDA-MB-231 were cultured in DMEM 1x (Gibco-Thermo Fisher Scientific, Waltham, Massachusetts, USA, Cat# 41966-029), supplemented with FBS 10% (BioWest, Cat# S1810-500) and L-glutamine (Gibco, Cat #25030-024). Same media was used for A549, further supplemented with non-essential amino acids (Gibco, Cat# 11140-035). HCT116 were cultured in McCoy’s (Gibco, Cat# 26600-023), supplemented with 10% FBS. We have induced brain tropism in these cell lines as previously described^12^. Briefly, GFP/Luc positive cells were injected intracardially in NSG mice and collected once brain metastases were established. After sorting the dissociated cells, these were cultured and re-injected intracardially in mice to generate cells lines with brain tropism (Br). We generated Br derivatives of MDA-fariaMB-231, A549 and HCT116, described from now on as MDA, A549 and HCT, respectively. All cell lines were genetically modified to overexpress UBE2C and MDA for the KD of UBE2C.

MET-CF78 is a patient-derived culture (PDC) established in our laboratory^26^ from a BM patient-derived xenograft (PDX). Briefly, cancer cells were isolated by enzymatic dissociation of a tumor derived from a subcutaneous PDX model which was implanted with BM sample from a patient with lung cancer. MET-CF78 were cultured in DMEM-F12 media (Gibco #11320-074) supplemented with 2% B-27, 1% N2 supplement, 1% L-glutamine (Gibco, Cat# 25030-024), 1x antibiotic-antimycotic (Gibco, Cat # 15240-096), rh-FGF (Stem Cell Technologies, Cat #02634) and rh-EGF (Sigma, Cat #E9644). Cells were seeded in previously coated flask with poly-L-ornithine and laminin.

### Cell modulation

To achieve the stable overexpression, the lentiviral vector was used for gene delivery. LeGO-iV2 (a gift from Boris Fehse, Addgene plasmid #27344) (Weber et al., 2008) was used to construct the recombinant lentiviral vector. The plasmid for human *UBE2C* overexpression was made by subcloning the PCR-amplified UBE2C (IDT) fragment into the EcoRI and Mscl (NEB) site of LeGO-iV2.

The lentiviral vector pLV hU6-sgRNA hUBC-dCas9-KRAB-T2A-Puro (a gift from Charles Gersbach, Addgene plasmid # 71236)^48^ was used for stable KD. sgRNAs were obtained from IDT. After annealing, sgRNAs were ligated to BsmBI digested pLV hU6-sgRNA hUBc-dCas9-KRAB-T2a-Puro vector. For lentiviral transfection, HEK293T cell lines were seeded in Petri dishes. Twenty-four hours after seeding, cells were incubated with lentiviral vector, helper, envelope plasmids, and polyethylenimine (PEI, Merck, Darmstadt, Germany, Cat# 408727) to increase the efficiency of infection. Cell culture medium containing the virus was collected 24 and 48 hours after infection. Target cells (MDA, A549 and HCT) were then infected with the produced virus using 2μg/ml polybrene (Merck, Darmstadt, Germany, #TR-1003-G). Stably transduced cells were either selected by flow cytometry using BD FACSAria III cell sorter or 1μg/ml puromycin (InvivoGen, San Diego, USA, #ant-pr-1).

### Immunoblotting

Whole cell lysates were prepared as previously described^49^. Briefly, cells were lysed in lysis buffer supplemented with 1x phosphatase inhibitors (PhosStop, Roche Diagnostics, Basel, Switzerland) and a 1x protease inhibitor cocktail (Complete Mini, Roche Diagnostics, Basel, Switzerland). After centrifugation at 10000g for 15 minutes at 4ºC, the supernatant was harvested. Total protein concentration was determined using the Bradford protein assay (BioRad, California, USA). Equal amounts of protein were subjected to sodium dodecyl sulfate–polyacrylamide gel electrophoresis and transferred onto nitrocellulose membranes (BioRad, California, USA), which were blocked with 5% skim milk for 1 hour at room temperature, incubated with specific primary antibodies overnight at 4ºC. Immunodetection was performed by incubation with horseradish-peroxidase–conjugated appropriate secondary antibodies and developed by chemiluminescence (Curix60, AGFA). The antibodies used were as follows: UBE2C (Boston Biochem, Cat# A650) and beta-actin (Abcam, Cat# ab8224). The latter was used as loading control. Two independent experiments were performed. The densitometry analysis was performed using Adobe Photoshop® software.

### Cell proliferation

Cell proliferation rate was determined at 24h, 48h, 72h and 96h using the CellTiter 96 Aqueous One Solution Reagent (MTS) (Promega, Wisconsin, USA, Cat# G3581) as defined by the manufacturer’s protocol. Briefly, in each time-point cells were plated in 96 well plates and incubated at 37ºC for 2h with MTS. The absorbance was measured at 490nm using the microplate reader Infinite M200 (Tecan, Crailsheim, Germany). Three independent experiments were performed with 3 technical replicates each.

### Colony Formation Assay

MDA or A549 single cell suspensions were seeded onto 6 well plates, with 100 cells per well, and incubated at 37ºC, 5% CO2. Cells were allowed to grow for two weeks in order to form single-cell-derived colonies. Wells were washed three times with PBS and fixed in 0.4% formaldehyde overnight and stained with 0.5% gentian violet for 2 minutes at room temperature. Staining was washed in distilled water, submerging the plates, and then allowed to air dry. Pictures of the plates were analyzed using ColonyArea plugin for ImageJ^50^, which calculates area and intensity of staining in colony formation assays.

### Migration

Cancer cells were seeded on the top chamber (2×10^4^ cells) of a CIM-Plate 16 (OMNI Life Science GmbH & Co KG, Bremen, Germany; Cat# 2801038) in triplicates. Cell migration was assessed in the xCELLigence system (ACEA Biosciences) in real time for 45 hours and readings were recorded every 15 minutes. Cells maintained in serum-free media served as a control.

### Invasion

Invasion assays were performed using 6.5mm Transwell® with 8.0μm pore size polyester membrane insert (Corning, New York, USA, Cat# 3464), coated with Matrigel® Matrix (Corning, New York, USA, Cat# 356237). Thirty thousand cancer cells were seeded on the top chamber in media with 5% FBS and the bottom chamber had 20% FBS media. Cells were allowed to migrate for 30h, at 37ºC and 5%CO_2_. After incubation, both chambers were carefully washed with PBS-Tween (0.05%), fixed with methanol (2 minutes at -20ºC) and stained with DAPI (Thermo Fisher Scientific, Waltham, Massachusetts, USA Cat# D1306) for 5 minutes at room temperature. Five random fields of each condition were acquired using Zeiss Axio Observer equipment (Zeiss, Jena, Germany), and the number of cells was counted manually in the Zen Blue software.

### *In vivo* orthotopic xenografts

In accordance with Directive 2010/63/EU (transposed to Portuguese legislation through Decreto-Lei No. 113/2013, of August 7th), all animal procedures were approved by the institutional animal welfare body (ORBEA-iMM), in order to ensure that the use of animals complies with all applicable legislation and following the 3R’s principle, as well as licensed by the Portuguese competent authority (license number: 012028\2016). All animals were kept in specific pathogen-free (SPF) conditions, randomly housed per groups under standard laboratory conditions (at 20-22°C under 10hour light/14hour dark), and given free access to food (RM3, SDS Diets, Witham, UK) and water (Ultrapure). Invasive procedures were performed with animals under anesthesia (ketamine, Ketamidor 100mg/ml 10ml, Plurivet; Medetomidina, Domtor 1mg/ml 10ml, Ecuphar), administered via intraperitoneal injection. Humane endpoints were established for 10% body weight loss, body condition scores ≤2 or lethargy, ataxia, bleeding, hunched/stiffed posture, self-mutilation, and skin bruising in consequence of tumor burden. Animals were euthanized using anesthetic overdose, using pentobarbital.

NSG mice were purchased from Charles River Laboratories (Massachusetts, USA) or obtained from a NSG colony established in-house. Animals were subjected to procedures between the ages of 11 and 22 weeks old. Cancer cells were injected intracranially in the frontal region of the cerebral cortex, using the bregma as reference point (2mm lateral right, 0mm anterior, and 2.5mm ventral). MDA cells were established by injecting 50000 cells, while 100000 MET-CF78 cells were injected in this model.

All mice were monitored for body weight, discomfort and distress every other day. Mice were euthanized, once any of the humane endpoints was reached. Histopathologic analysis was performed in CNS samples.

### *In vivo* imaging of mice

Mice were imaged under anesthesia. Prior to the image acquisition, animals were injected by intraperitoneal injection with XenoLight D-Luciferin, Potassium Salt (Perkin Elmer, Boston, USA, Cat# PELS122799) subcutaneously. After 10 minutes, bioluminescence images were acquired using IVIS Lumina System using 5 minutes exposure (Perkin Elmer), and analyzed using Living Image software, version 3.0.

### Histopathological analysis of mouse samples

Tissue samples were fixed immediately in 10% neutral buffered formalin solution, dehydrated, and embedded in paraffin, serially sectioned at a thickness of 5μm using a microtome, mounted on microscope slides and stained with hematoxylin and eosin (H&E) for morphological examination. H&E slides were blindly examined by two independent researchers and a specialized pathologist, and representative photomicrographs were taken using the NanoZoomer SQ slide scanner (Hamamatsu Photonics, Hamamatsu City, Japan) with 20x magnification (0.46μm resolution).

### Drug screening

Drug screening was conducted using Corning 384-well microtiter plates pre-dispensed with inhibitors dissolved in DMSO prior to cell seeding using D300e Drug Dispenser (Tecan, Crailsheim, Germany), sealed using Parafilm and stored at -80°C until use. Stock solutions for printing were prepared at 10mM concentration. A series of six-nine dilution steps of each inhibitor between 32.5 and 25000nM was printed in logarithmic distribution and DMSO content was normalized to 0.25% in all wells. Staurosporine was added to each plate as a positive control. DMSO and empty wells served as negative controls. In addition, to avoid plate effects, the two outer columns and rows were not used, and the inner part was dispensed in a randomized fashion. One hour before testing, the assay plates were removed from -80°C and thawed at room temperature.

To improve screening stability, all (MDA and HCT) cell lines underwent a cell density optimization procedure. Using MultiDrop Combi Reagent Dispenser (Thermo Fisher Scientific, Schwerte, Germany), 30μl of cell suspension were seeded into each well of the pre-printed plates. After 72 hours of incubation at 37ºC and 5% CO_2_, plates were taken out of the incubator and equilibrated to room temperature for 30 minutes. Thirty microliters of CellTiter-Glo reagent (Promega GmbH - Walldorf, Germany Cat# G7573) was added to each well and plates were incubated 10 minutes at 37ºC and posteriorly read in the Spark MultiMode Plate reader (Tecan, Crailsheim, Germany).

### *In vitro* drug testing

*In vitro* drug assays were performed by seeding MDA, A549 (1000 cells) or MET-CF78 (500 cells) in 96-well plates and using different concentrations (25nM, 100nM, 250nM, 500nM and 1mM) of dactolisib (Selleckchem, Munich, Germany, Cat# S1009) and Genz-644282 (Selleckchem, Munich, Germany, Cat# S0093). Proliferation assays were performed as described above.

### *In vivo* drug testing using orthotopic BM xenografts

*In vivo* drug response evaluation was performed using NSG mice injected intracranially with MDA cancer cell line (5×10^5^ cells/mouse). Dual ATP-competitive PI3K and mTOR inhibitor Dactolisib (30mg/kg) was purchased from Selleck Chemicals (Munich, Germany). Dactolisib was freshly solved in N-Methyl-2-Pyrrolidone (NMP; Sigma-Aldrich, Darmstadt, Germany, Cat# 328634) and Polyethylene glycol 300 (PEG300; Sigma-Aldrich, Darmstadt, Germany, Cat# 202371) (10/90, v/v) immediately before administration by oral gavage, weight-adjusted. Animals were randomized into one treatment group and one untreated control group (NMP+PEG300). Treatment started four- or seven-days post-injection and mice received two cycles of therapy (five days on and two days off), until day 13 or 17 post-injection (**Fig. 6A** and **H**). The animals were monitored daily, and body weight variations were recorded throughout treatment. Mice were euthanized by the end of the treatment (day 15 or 18), and CNS and organs (lungs, liver, spleen and kidneys) were collected for histopathological analysis, as described above.

### UBE2C IHC in orthotopic BM xenografts samples

In mice injected with MDA and treated with dactolisib (or vehicle), UBE2C downregulation was analyzed by an in-house developed macro for ImageJ/Fiji. Images of the intracranial tumors were acquired using the Nanozoomer SQ system (acquisition: 20x; export: 40x magnification) and analyzed using the ImageJ/Fiji macro to quantify the percentage of high UBE2C-staining in the tumor tissue (available at https://github.com/ClaraBarreto/UBE2C). Briefly, the total tumorous tissue area (TTA) is determined by using the “Color Threshold” feature in the HSB color space, after manual exclusion of non-tumorous tissue and staining artifacts regions. Two classes of positive staining were then defined: dark brown for strong staining and light brown for weak staining. Threshold values for the “Color Threshold” feature were then empirically determined for the dark-brown area (DBA) using the RGB color space, by analyzing a set of representative images with strong staining. The light-brown area (LBA) was then calculated using the formula: LBA = TTA – DBA, and area fractions were determined as follows: %LBA = LBA/TTA and %DBA = DBA/TTA.

### Statistical analysis

Statistical differences were determined using t-test (parametric) or Mann– Whitney tests (non-parametric) on GraphPad Prism v6.0 (GraphPad, California, USA, GraphPad Prism, RRID:SCR_002798), as stated in figure legends Differences were considered statistically significant for p≤0.05.

## AKNOWLEDGMENTS

The authors would like to acknowledge the patients who kindly provided the tumor specimens included in the RNA sequencing and TMA analysis for this research, and the Biobanco-iMM CAML who enabled the process of tumor specimen collection, processing and storage. Finally, the authors thank Pedro Pereira, the Histology and Comparative Pathology Laboratory and the Rodent Facility from Instituto de Medicina Molecular João Lobo Antunes for technical assistance.

## AUTHOR CONTRIBUTIONS

Study design: EP, RC, JTB and CCF. Study conduct: EP, RC, CC, NQ, DPicard, DPauck, TC, PR, RR, CF, JP and CCF. Data collection: EP, RC, CC, NQ, DPicard, DPauck, DD and JC. Data analysis: EP, RC, CC, NQ, DPicard, DPauck, TC, PR, CB, and CCF. Data interpretation: EP, RC, DPicard, TC, PR, JTB and CCF. Drafting manuscript: EP, RC and CCF. Revising manuscript content: EP, RC, CC, NQ, DPicard, DPauck, TC, PR, CB, DD, JC, RR, JP, JM, MR, JTB and CCF. All authors approved the final version of this manuscript and take responsibility for the integrity of the data analysis.

## REFERENCES

1 Kocher, M. et al. Adjuvant whole-brain radiotherapy versus observation after radiosurgery or surgical resection of one to three cerebral metastases: results of the EORTC 22952-26001 study. J Clin Oncol 29, 134–141, doi:10.1200/JCO.2010.30.1655 (2011).

2 Berghoff, A. S. et al. Descriptive statistical analysis of a real life cohort of 2419 patients with brain metastases of solid cancers. ESMO Open 1, e000024, doi:10.1136/esmoopen-2015-000024 (2016).

3 Achrol, A. S. et al. Brain metastases. Nat Rev Dis Primers 5, 5, doi:10.1038/s41572-018-0055-y (2019).

4 Nayak, L., Lee, E. Q. & Wen, P. Y. Epidemiology of brain metastases. Curr Oncol Rep 14, 48–54, doi:10.1007/s11912-011-0203-y (2012).

5 Barnholtz-Sloan, J. S. et al. Incidence proportions of brain metastases in patients diagnosed (1973 to 2001) in the Metropolitan Detroit Cancer Surveillance System. J Clin Oncol 22, 2865–2872, doi:10.1200/JCO.2004.12.149 (2004).

6 Smedby, K. E., Brandt, L., Backlund, M. L. & Blomqvist, P. Brain metastases admissions in Sweden between 1987 and 2006. Br J Cancer 101, 1919–1924, doi:10.1038/sj.bjc.6605373 (2009).

7 Rosenfelder, N. & Brada, M. Integrated treatment of brain metastases. Curr Opin Oncol 31, 501–507, doi:10.1097/CCO.0000000000000573 (2019).

8 Beauchesne, P. Intrathecal chemotherapy for treatment of leptomeningeal dissemination of metastatic tumours. The Lancet Oncology 11, 871–879, doi:10.1016/s1470-2045(10)70034-6 (2010).

9 Fortin, D. The blood-brain barrier: its influence in the treatment of brain tumors metastases. Curr Cancer Drug Targets 12, 247–259, doi:10.2174/156800912799277511 (2012).

10 Brastianos, P. K. et al. Genomic Characterization of Brain Metastases Reveals Branched Evolution and Potential Therapeutic Targets. Cancer Discov 5, 1164–1177, doi:10.1158/2159-8290.CD-15-0369 (2015).

11 Eichler, A. F. et al. The biology of brain metastases-translation to new therapies. Nat Rev Clin Oncol 8, 344–356, doi:10.1038/nrclinonc.2011.58 (2011).

12 Bos, P. D. et al. Genes that mediate breast cancer metastasis to the brain. Nature 459, 1005–1009, doi:10.1038/nature08021 (2009).

13 Valiente, M. et al. Serpins promote cancer cell survival and vascular co-option in brain metastasis. Cell 156, 1002–1016, doi:10.1016/j.cell.2014.01.040 (2014).

14 Townsley, F. M., Aristarkhov, A., Beck, S., Hershko, A. & Ruderman, J. V. Dominant-negative cyclin-selective ubiquitin carrier protein E2-C/UbcH10 blocks cells in metaphase. Proc Natl Acad Sci U S A 94, 2362–2367, doi:10.1073/pnas.94.6.2362 (1997).

15 Reddy, S. K., Rape, M., Margansky, W. A. & Kirschner, M. W. Ubiquitination by the anaphase-promoting complex drives spindle checkpoint inactivation. Nature 446, 921–925, doi:10.1038/nature05734 (2007).

16 Okamoto, Y. et al. UbcH10 is the cancer-related E2 ubiquitin-conjugating enzyme. Cancer Res 63, 4167–4173 (2003).

17 Wagner, K. W. et al. Overexpression, genomic amplification and therapeutic potential of inhibiting the UbcH10 ubiquitin conjugase in human carcinomas of diverse anatomic origin. Oncogene 23, 6621–6629, doi:10.1038/sj.onc.1207861 (2004).

18 Sjodahl, G. et al. A molecular taxonomy for urothelial carcinoma. Clin Cancer Res 18, 3377–3386, doi:10.1158/1078-0432.CCR-12-0077-T (2012).

19 Pawitan, Y. et al. Gene expression profiling spares early breast cancer patients from adjuvant therapy: derived and validated in two population-based cohorts. Breast Cancer Res 7, R953–964, doi:10.1186/bcr1325 (2005).

20 Bogunovic, D. et al. Immune profile and mitotic index of metastatic melanoma lesions enhance clinical staging in predicting patient survival. Proc Natl Acad Sci U S A 106, 20429–20434, doi:10.1073/pnas.0905139106 (2009).

21 Frei, E. et al. Addition of rituximab to chemotherapy overcomes the negative prognostic impact of cyclin E expression in diffuse large B-cell lymphoma. J Clin Pathol 66, 956–961, doi:10.1136/jclinpath-2013-201619 (2013).

22 Hanamura, I., Huang, Y., Zhan, F., Barlogie, B. & Shaughnessy, J. Prognostic value of cyclin D2 mRNA expression in newly diagnosed multiple myeloma treated with high-dose chemotherapy and tandem autologous stem cell transplantations. Leukemia 20, 1288–1290, doi:10.1038/sj.leu.2404253 (2006).

23 Su, Z. et al. An investigation of biomarkers derived from legacy microarray data for their utility in the RNA-seq era. Genome Biol 15, 523, doi:10.1186/s13059-014-0523-y (2014).

24 Yang, S. et al. A Novel MIF Signaling Pathway Drives the Malignant Character of Pancreatic Cancer by Targeting NR3C2. Cancer Res 76, 3838–3850, doi:10.1158/0008-5472.CAN-15-2841 (2016).

25 Gonzalez, H. et al. Cellular architecture of human brain metastases. Cell 185, 729–745 e720, doi:10.1016/j.cell.2021.12.043 (2022).

26 Faria, C. C. et al. Patient-derived models of brain metastases recapitulate human disseminated disease. Cell Rep Med 3, 100623, doi:10.1016/j.xcrm.2022.100623 (2022).

27 Gil del Alcazar, C. R. et al. Inhibition of DNA double-strand break repair by the dual PI3K/mTOR inhibitor NVP-BEZ235 as a strategy for radiosensitization of glioblastoma. Clin Cancer Res 20, 1235–1248, doi:10.1158/1078-0432.CCR-13-1607 (2014).

28 Perrotta, I. et al. Immunohistochemical analysis of the ubiquitin-conjugating enzyme UbcH10 in lung cancer: a useful tool for diagnosis and therapy. J Histochem Cytochem 60, 359–365, doi:10.1369/0022155412439717 (2012).

29 Guo, J. et al. Deregulation of UBE2C-mediated autophagy repression aggravates NSCLC progression. Oncogenesis 7, 49, doi:10.1038/s41389-018-0054-6 (2018).

30 Loussouarn, D. et al. Validation of UBE2C protein as a prognostic marker in node-positive breast cancer. Br J Cancer 101, 166–173, doi:10.1038/sj.bjc.6605122 (2009).

31 Jayanthi, V., Das, A. B. & Saxena, U. Grade-specific diagnostic and prognostic biomarkers in breast cancer. Genomics 112, 388–396, doi:10.1016/j.ygeno.2019.03.001 (2020).

32 Wang, C. et al. Knockdown of UbcH10 enhances the chemosensitivity of dual drug resistant breast cancer cells to epirubicin and docetaxel. Int J Mol Sci 16, 4698–4712, doi:10.3390/ijms16034698 (2015).

33 Presta, I. et al. UbcH10 a Major Actor in Cancerogenesis and a Potential Tool for Diagnosis and Therapy. Int J Mol Sci 21, doi:10.3390/ijms21062041 (2020).

34 Fujita, T. et al. Overexpression of UbcH10 alternates the cell cycle profile and accelerate the tumor proliferation in colon cancer. BMC Cancer 9, 87, doi:10.1186/1471-2407-9-87 (2009).

35 Berlingieri, M. T. et al. UbcH10 expression may be a useful tool in the prognosis of ovarian carcinomas. Oncogene 26, 2136–2140, doi:10.1038/sj.onc.1210010 (2007).

36 Morikawa, T. et al. UBE2C is a marker of unfavorable prognosis in bladder cancer after radical cystectomy. Int J Clin Exp Pathol 6, 1367–1374 (2013).

37 Liu, Y. et al. UBE2C Is Upregulated by Estrogen and Promotes Epithelial-Mesenchymal Transition via p53 in Endometrial Cancer. Mol Cancer Res 18, 204–215, doi:10.1158/1541-7786.MCR-19-0561 (2020).

38 Wang, R. et al. UBE2C induces EMT through Wnt/betacatenin and PI3K/Akt signaling pathways by regulating phosphorylation levels of Aurora-A. Int J Oncol 50, 1116–1126, doi:10.3892/ijo.2017.3880 (2017).

39 Xiong, Y. et al. UBE2C functions as a potential oncogene by enhancing cell proliferation, migration, invasion, and drug resistance in hepatocellular carcinoma cells. Biosci Rep 39, doi:10.1042/BSR20182384 (2019).

40 Venur, V. A., Chukwueke, U. N. & Lee, E. Q. Advances in Management of Brain and Leptomeningeal Metastases. Curr Neurol Neurosci Rep 20, 26, doi:10.1007/s11910-020-01039-1 (2020).

41 Scott, B. J. & Kesari, S. Leptomeningeal metastases in breast cancer. Am J Cancer Res 3, 117–126 (2013).

42 Le Rhun, E. et al. EANO-ESMO Clinical Practice Guidelines for diagnosis, treatment and follow-up of patients with leptomeningeal metastasis from solid tumours. Ann Oncol 28, iv84–iv99, doi:10.1093/annonc/mdx221 (2017).

43 Yang, J. C. H. et al. Osimertinib in Patients With Epidermal Growth Factor Receptor Mutation-Positive Non-Small-Cell Lung Cancer and Leptomeningeal Metastases: The BLOOM Study. J Clin Oncol 38, 538–547, doi:10.1200/JCO.19.00457 (2020).

44 Brastianos, P. K. et al. Phase II study of pembrolizumab in leptomeningeal carcinomatosis. Journal of Clinical Oncology 36, 2007–2007, doi:10.1200/JCO.2018.36.15_suppl.2007 (2018).

45 Smalley, I. et al. Proteomic Analysis of CSF from Patients with Leptomeningeal Melanoma Metastases Identifies Signatures Associated with Disease Progression and Therapeutic Resistance. Clin Cancer Res 26, 2163–2175, doi:10.1158/1078-0432.CCR-19-2840 (2020).

46 Wang, H. et al. CCI-779 inhibits cell-cycle G2-M progression and invasion of castration-resistant prostate cancer via attenuation of UBE2C transcription and mRNA stability. Cancer Res 71, 4866–4876, doi:10.1158/0008-5472.CAN-10-4576 (2011).

47 Sloan, C. A. et al. ENCODE data at the ENCODE portal. Nucleic Acids Res 44, D726–732, doi:10.1093/nar/gkv1160 (2016).

48 Thakore, P. I. et al. Highly specific epigenome editing by CRISPR-Cas9 repressors for silencing of distal regulatory elements. Nat Methods 12, 1143–1149, doi:10.1038/nmeth.3630 (2015).

49 Barata, J. T. et al. Activation of PI3K is indispensable for interleukin 7-mediated viability, proliferation, glucose use, and growth of T cell acute lymphoblastic leukemia cells. J Exp Med 200, 659–669, doi:10.1084/jem.20040789 (2004).

50 Guzman, C., Bagga, M., Kaur, A., Westermarck, J. & Abankwa, D. ColonyArea: an ImageJ plugin to automatically quantify colony formation in clonogenic assays. PLoS One 9, e92444, doi:10.1371/journal.pone.0092444 (2014).

