## Supplementary Materials for "UBE2C promotes leptomeningeal dissemination and is a therapeutic target in brain metastatic disease"

**A**

| Tissue of origin | Microarray datasets (number of samples) GSE2109 (Tumor) | GSE7307 (Normal) |
|---|---|---|
| **Bladder** | 30 | 0 |
| **Brain** | 7 | 114 |
| **Breast** | 354 | 9 |
| **Colon** | 381 | 0 |
| **Esophagus** | 7 | 4 |
| **Lung** | 132 | 0 |
| **Skin** | 1 | 7 |
| **Uterus** | 240 | 45 |
| **Unknown** | - | - |

**B**

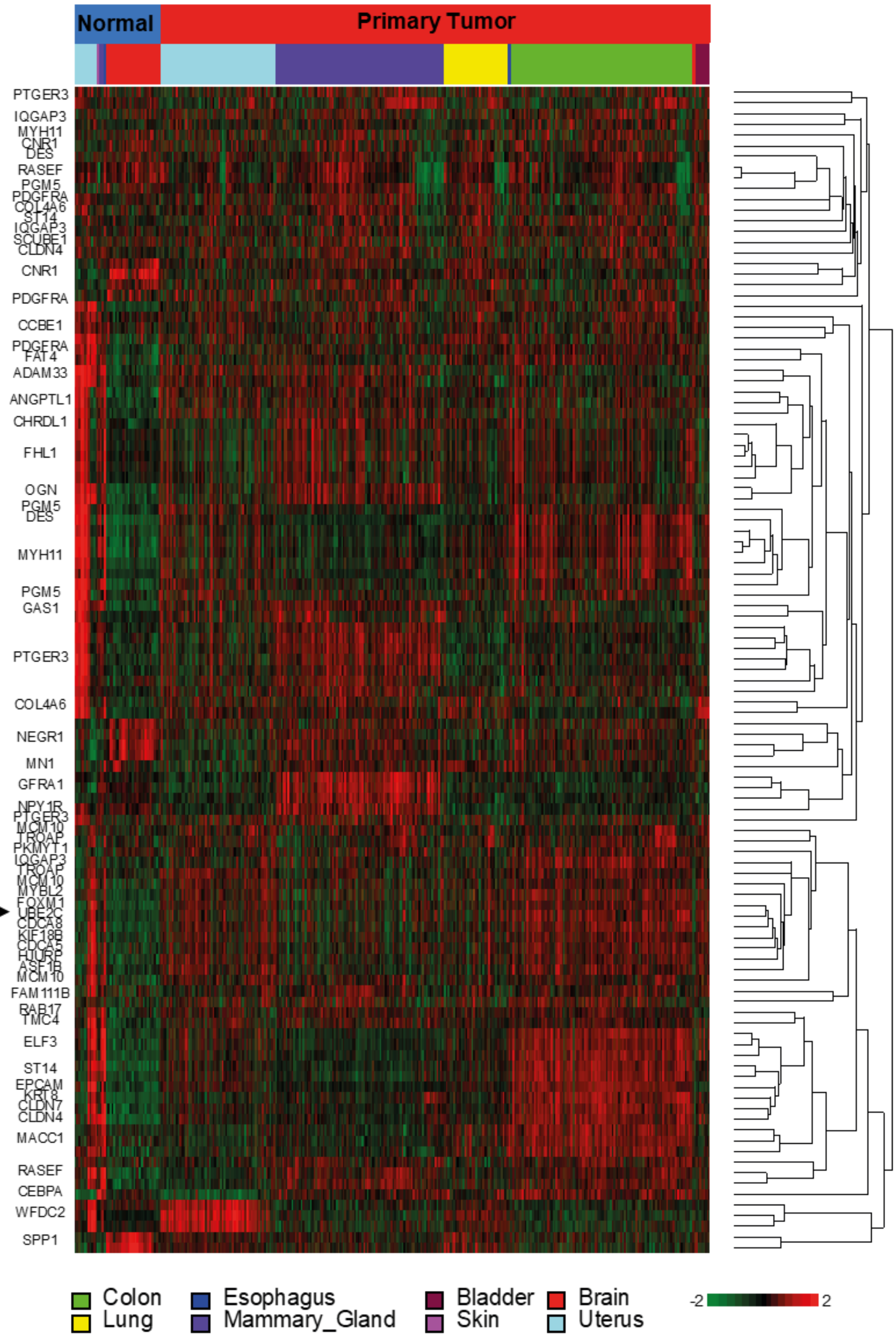

**Supplemental Figure 1. Genes differentially expressed between diverse primary tumors and normal tissue samples. A.** Datasets included in the microarray bioinformatic analysis, including data from primary tumors and normal tissue samples. **B.** Heatmap representing the expression of the top upregulated genes in the microarray data of primary tumors and normal tissue samples.

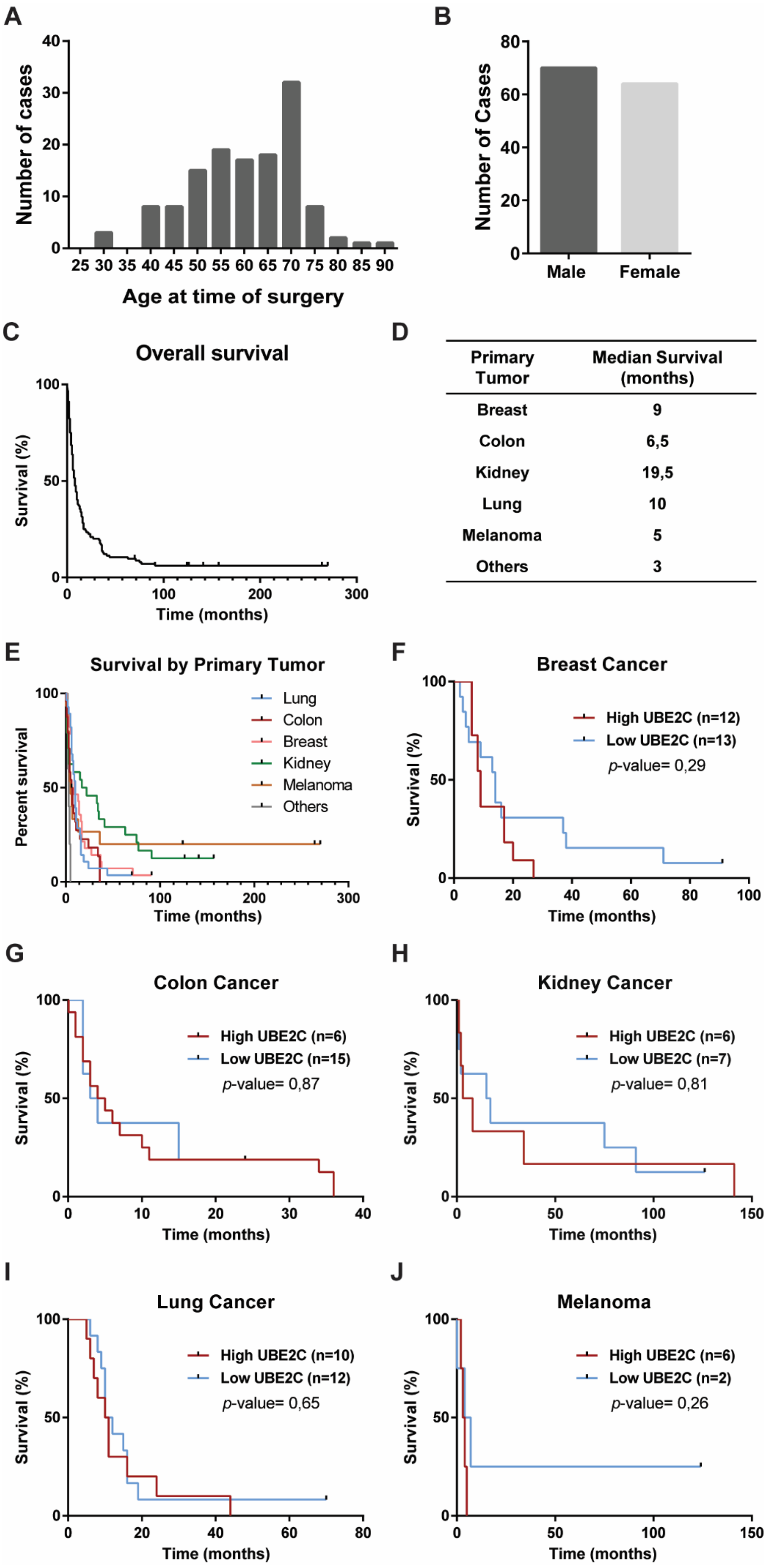


| Primary Tumor | Median Survival (months) |
|---|---|
| Breast | 9 |
| Colon | 6,5 |
| Kidney | 19,5 |
| Lung | 10 |
| Melanoma | 5 |
| Others | 3 |

**Supplemental Figure 2. Characterization of the cohort of patients with BM used in tissue microarrays (validation cohort). A.** Age distribution. **B.** Gender distribution. **C.** Overall survival of patients since the diagnosis of BM. **D.** Median survival of patients with BM according to the primary tumor origin. **E.** Kaplan-Meier analysis of patients' survival according to the UBE2C protein levels (high vs low) in BM patients with **(F)** breast, **(G)** colon, **(H)** kidney, **(I)** lung cancer, and **(J)** melanoma. According to the Log-rank (Mantel-Cox) test, differences were considered statistically significant for *p*-values≤0.05.

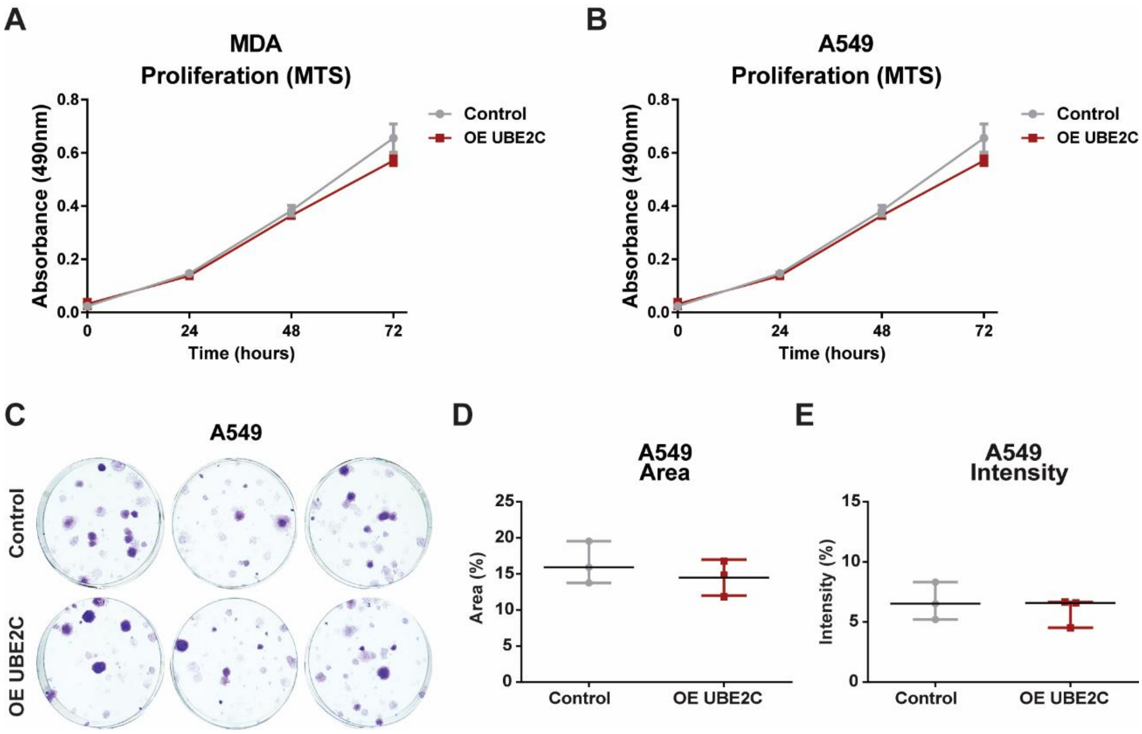


**Supplemental Figure 3.** ***In vitro*** **proliferation of MDA and A549 cell lines with** ***UBE2C*** **overexpression.** MTS assay to assess proliferation in cancer cell lines with overexpression of *UBE2C* in (**A**) MDA (breast cancer; n=2) and (**B**) A549 (lung cancer; n=1). **(C)** Representative pictures of colony formation assays (CFA) performed using A549 OE *UBE2C* (100 cells/well) with **(D)** quantification of the area, *p*-value=0.44, and **(E)** intensity, *p*-value=0.54. CFA was quantified using the plugin ColonyArea on ImageJ. Mann-Whitney test. Data is represented as median with an interquartile range.

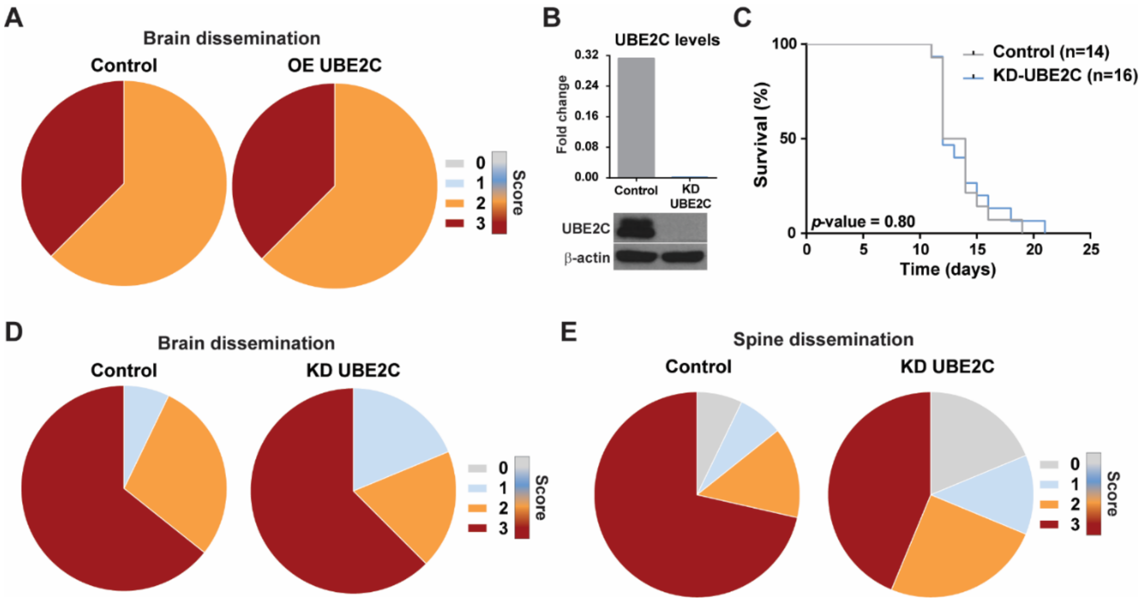


**Supplemental Figure 4. Orthotopic mouse models injected with breast cancer cells with KD *UBE2C*. A.** Brain dissemination score of animals injected with MDA control cells or OE of *UBE2C* (n=8/group). Both groups presented the same scores (2: 62,5%, 3: 37,5%). **B.** Western Blot of UBE2C levels in MDA cells with knockdown (KD) of *UBE2C* and respective quantification by densitometry analysis. **C.** Kaplan-Meier analysis of survival in orthotopic xenografts (control: n=14; KD *UBE2C*: n=16). **D.** Percentage of brain dissemination in control (1: 7,143%, 2: 28,571, 3: 64,286%) and KD *UBE2C* (1: 18,75%, 2: 18,75%, 3: 62,5%) groups. **E.** Percentage of spine dissemination in control (0: 7,143%, 1: 7,143%, 2: 14,286%, 3: 71,429%) or KD *UBE2C* (0: 18,75%, 1: 12,5%, 2: 25%, 3: 43,75%) animals. Score used to assess the leptomeningeal dissemination: 0- negative; 1- mild; 2- moderate; 3- marked

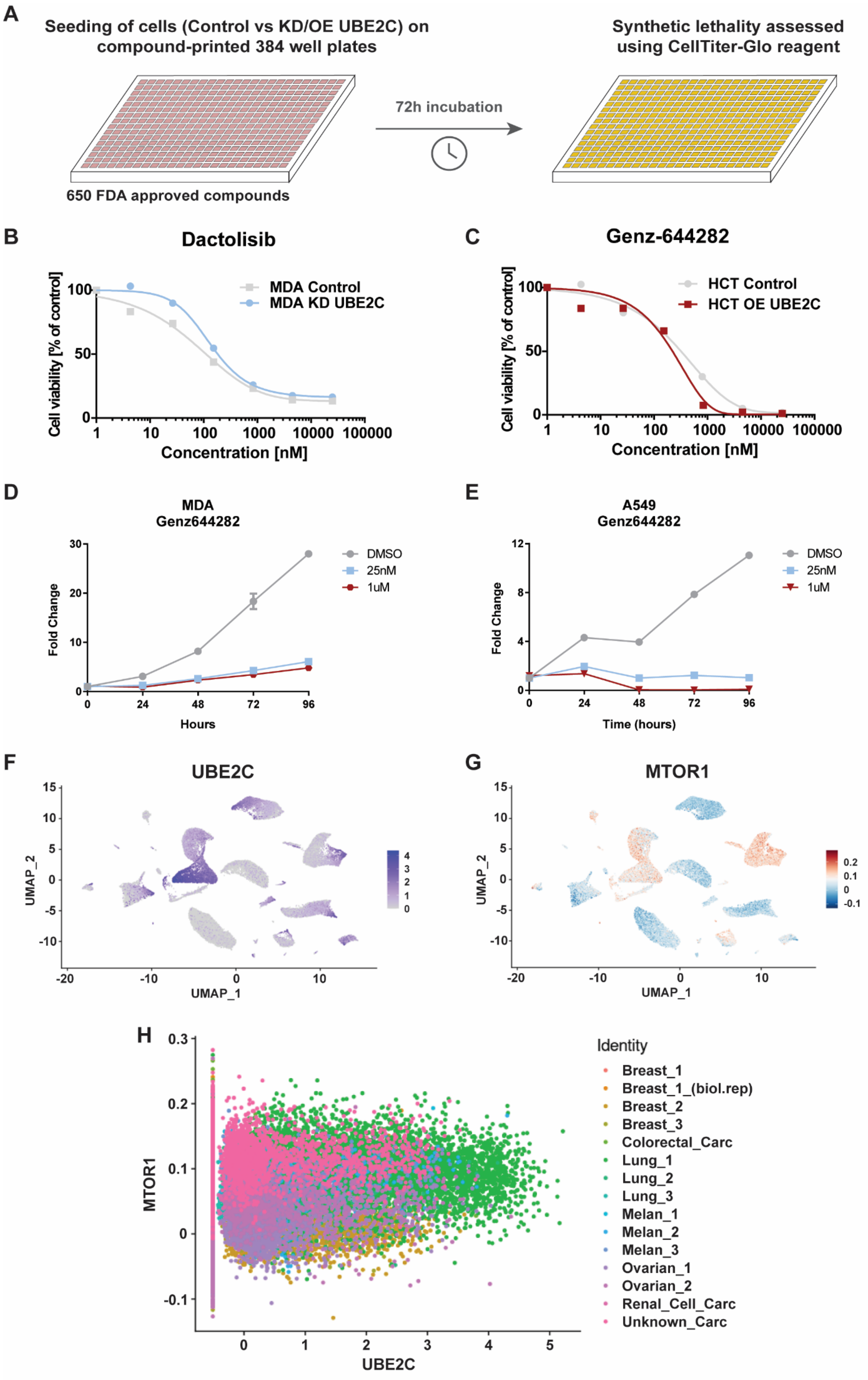

A
Seeding of cells (Control vs KD/OE UBE2C) on compound-printed 384 well plates
72h incubation
Synthetic lethality assessed using CellTiter-Glo reagent
650 FDA approved compounds
B
Dactolisib
MDA Control
MDA KD UBE2C
Cell viability [% of control]
Concentration [nM]
C
Genz-644282
HCT Control
HCT OE UBE2C
Cell viability [% of control]
Concentration [nM]
D
MDA
Genz644282
DMSO
25nM
1uM
Fold Change
Hours
E
A549
Genz644282
DMSO
25nM
1uM
Fold Change
Time (hours)
F
UBE2C
UMAP_2
UMAP_1
G
MTOR1
UMAP_2
UMAP_1
H
Identity
Breast_1
Breast_1_(biol.rep)
Breast_2
Breast_3
Colorectal_Carc
Lung_1
Lung_2
Lung_3
Melan_1
Melan_2
Melan_3
Ovarian_1
Ovarian_2
Renal_Cell_Carc
Unknown_Carc
MTOR1
UBE2C

**Supplemental Figure 5. Dactolisib and Genz644282 are potential inhibitors of *UBE2C*-expressing cells. A.** Drug screening performed by seeding cells on 384-well plates pre-dispensed with inhibitors dissolved in DMSO (6-9 dilution steps for each inhibitor ranging from 32.5-25000nM). After 72 hours, treated cells were incubated with CellTiter-Glo reagent, and absorbance was used as a readout of cell viability. **B.** Effect of PI3K/mTOR inhibition (dactolisib) in breast cancer cell line MDA (control vs KD *UBE2C*). **C.** Effect of topoisomerase I inhibition (Genz644282) in a colon cancer cell line HCT (control vs OE *UBE2C*). A four-parameter logistic dose-response curve was used to describe the association between response to treatment and drug concentration. MTS assays using 25nM and 1μM concentrations of Genz644282 to assess proliferation in (**D**) MDA and (**E**) A459 cell lines with OE of *UBE2C* Metastatic tumor cells (MTCs) visualized in a UMAP plot of GSE186344. Cells were annotated by mRNA expression of **(F)** UBE2C (purple is high, and grey is no expression) or by **(G)** geneset enrichment score of HALLMARK_PI3K_AKT_MTOR_SIGNALING (red is high and blue is low) in 15 metastatic samples. **H.** Scatter plot of mRNA expression and geneset enrichment scores shows a positive correlation between UBE2C and MTOR1 (r=0,21; *p*-value= $2.2x10^{-16}$); data from Morikawa *et al.*, 2022.

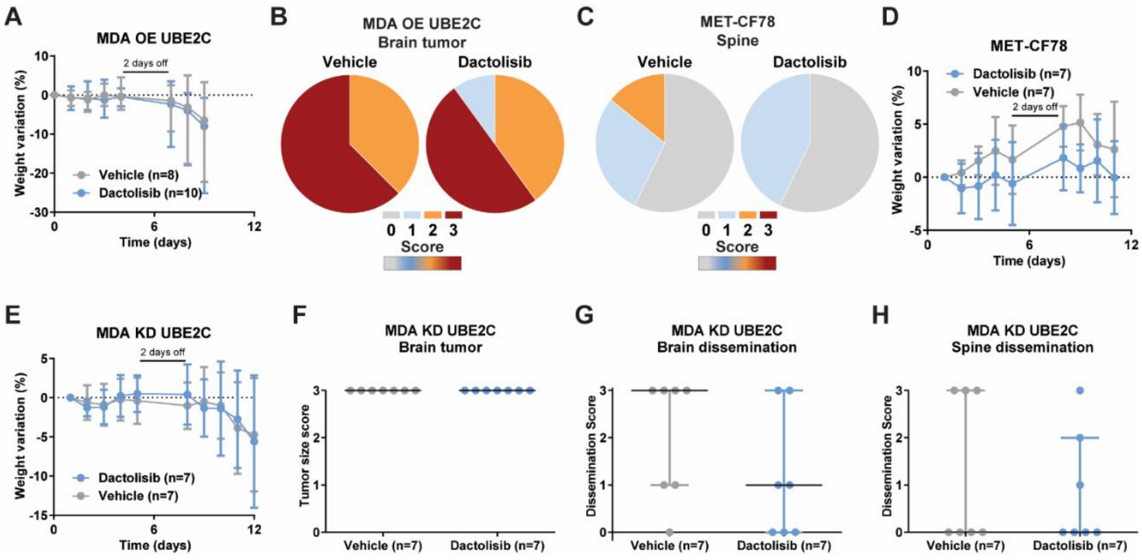


**Supplemental Figure 6. Dactolisib treatment of orthotopic xenografts with *UBE2C* modulation and patient-derived xenografts (MET-CF 78). A.** Weight variation of NSG mice injected intracranially with MDA OE *UBE2C* and treated with dactolisib (n=10) or vehicle (n=8). **B.** Percentage of brain tumor dissemination in MDA OE *UBE2C* mice treated with vehicle (2: 37,5%, 3: 62,5%) or dactolisib (1: 10%, 2: 40%, 3: 50%). **C.** Percentage of spine dissemination in patient-derived xenografts from a lung cancer BM (MET-CF78) treated with vehicle (0: 57,143; 2: 28,571%, 3: 14,286%) or dactolisib (0: 57,143%, 1: 42,857%); (n=7/group). **D.** Weight variation of treated PDXs. **E.** Weight variation of MDA KD *UBE2C* mice treated with vehicle or dactolisib (n=7/group). Histopathological scoring of **(F)** brain tumor size (0- no tumor; 1- minimal to mild; 2- moderate; 3- marked), p>0,999, and leptomeningeal dissemination **(G)** in the brain, p=0.347; and **(H)** in the spine, p=0,755 (Score used to assess the leptomeningeal dissemination: 0- negative; 1- mild; 2- moderate; 3- marked.); Mann-Whitney test. Data represented as median with interquartile range.
